# Unraveling Morphogenesis, Starvation, and Light Responses in a Mushroom-Forming Fungus, *Coprinopsis cinerea*, Using Long Read Sequencing and Extensive Expression Profiling

**DOI:** 10.1101/2024.05.10.593147

**Authors:** Botond Hegedüs, Neha Sahu, Balázs Bálint, Sajeet Haridas, Viktória Bense, Zsolt Merényi, Máté Virágh, Hongli Wu, Xiao-Bin Liu, Robert Riley, Anna Lipzen, Maxim Koriabine, Emily Savage, Jie Guo, Kerrie Barry, Vivian Ng, Péter Urbán, Attila Gyenesei, Michael Freitag, Igor V. Grigoriev, László G. Nagy

**Affiliations:** Synthetic and Systems Biology Unit, Institute of Biochemistry, HUN-REN Biological Research Center, Temesvári krt. 62, Szeged H-6726, Hungary; U.S. Department of Energy Joint Genome Institute, Lawrence Berkeley National Laboratory, Berkeley, CA 94720, United States of America; János Szentágothai Research Center, University of Pécs, Ifjúság útja 20, Pécs H-7624, Hungary; Department of Biochemistry and Biophysics, Oregon State University, Corvallis, OR 97331, United States of America; Department of Plant and Microbial Biology, University of California Berkeley, Berkeley, CA 94720, United States of America

## Abstract

Mushroom-forming fungi (Agaricomycetes) are emerging as pivotal players in several fields, as drivers of nutrient cycling, sources of novel applications, and the group includes some of the most morphologically complex multicellular fungi. Genomic data for Agaricomycetes are accumulating at a steady pace, however, this is not paralleled by improvements in the quality of genome sequence and associated functional gene annotations, which leaves gene function notoriously poorly understood in comparison with other fungi and model eukaryotes. We set out to improve our functional understanding of the model mushroom *Coprinopsis cinerea* by integrating a new, chromosome-level assembly with high-quality gene predictions and functional information derived from gene-expression profiling data across 67 developmental, stress, and light conditions. The new annotation has considerably improved quality metrics and includes 5’- and 3’-untranslated regions (UTRs), polyadenylation sites (PAS), upstream ORFs (uORFs), splicing isoforms, conserved sequence motifs (e.g., TATA and Kozak boxes) and microexons. We found that alternative polyadenylation is widespread in *C. cinerea*, but that it is not specifically regulated across the various conditions used here. Transcriptome profiling allowed us to delineate core gene sets corresponding to carbon starvation, light-response, and hyphal differentiation, and uncover new aspects of the light-regulated phases of life cycle. As a result, the genome of *C. cinerea* has now become the most comprehensively annotated genome among mushroom-forming fungi, which will contribute to multiple rapidly expanding fields, including research on their life history, light and stress responses, as well as multicellular development.

## Introduction

Global concern for sustainability increased interest in environmentally friendly solutions for old industrial problems, for which the kingdom fungi offers remarkable biological agents. The class Agaricomycetes (mushroom-forming fungi) contains the most powerful degraders of plant lignocellulosic biomass and species that produce the most morphologically complex fruiting bodies, which are commercially cultivated by the rapidly expanding mushroom industry(1, 2). These two traits, among others, have fueled Agaricomycete genomics, which led to the proliferation of draft genome sequences, of which currently there are >500 in various repositories, such as MycoCosm (3). However, a limiting factor in documenting the genetics of key traits in the Agaricomycetes, as compared to the Ascomycota, has been the paucity of high quality annotations on chromosome-level genome assemblies, and the shortage of knowledge on gene function. This stems from the large diversity of basidiomycete model systems (4), the lack of systematic gene deletion projects, and the difficulty of extrapolating gene function from well-researched organisms, such as *Saccharomyces cerevisiae*, simply based on homology. Recent efforts have culminated in the publication of several chromosome-level assemblies of various Agaricomycetes (5–11). While these genomes represent a significant step in understanding the biology of these organisms, improvements of gene annotations did not follow that of contiguity. To the best of our knowledge, there is no Agaricomycetes species in which full-length transcripts, untranslated regions (UTRs), intergenic regions, upstream ORFs (uORFs) or polyadenylation sites (PAS) have been determined with high precision. The problem cannot be easily alleviated by *in silico* predictions due to the genome architecture of fungi, necessitating the use of appropriate experimental data for precise annotations.

Fungal genomes are relatively compact, usually with short intergenic spaces and few and short introns (12). Gene spacing is often so tight that the untranslated regions (UTRs) of adjacent genes overlap. Overlaps of 3’ UTRs of adjacent convergently transcribed genes are widespread in fungi, and can play direct or indirect roles in regulating each other’s expression (13, 14). The situation is further complicated by prevalent polycistronic transcription (15, 16), which could be the result from transcriptional readthrough due to weak termination signals (15). The identification of the UTR regions is important because they can contain (un)structured elements, such as uORFs in the 5’ UTR that may modulate translation initiation and stalling (17–19). The 3’ UTR may contain elements that regulate mRNA stability, translation, or localization (19). The presence of functional elements on the 3’UTR region is regulated through the process of alternative polyadenylation (APA) (20–22).

*Coprinopsis cinerea* is one of the most widely used model mushroom species, with a history of investigations dating back to the early 20th century (23). It has been used to understand principles of mushroom development, meiosis and mitosis, and photobiology, among others, and is currently one of the main model species for research on fungal multicellularity (24) and developmental biology (23, 25, 26). In terms of lifestyle, *C. cinerea* is a litter decomposer (LD), it feeds on non-woody lignocellulosic plant biomass, such as grasses, straw or manure (27). Two chromosome-level and a draft assembly for the two most relevant strains (Okayama7 #130 and Amut1Bmut1 #326) have been published (26, 28, 29). This fungus is easy to culture in the laboratory on defined media (30). In wildtype strains haploid spores (1n) germinate into monokaryotic hyphae (1n), which fuse to form dikaryons (2n), which can produce sexual fruiting bodies and spores, if different alleles are present at both *A* and *B* mating-type loci. In contrast, the widely used Amut1Bmut1 #326 strain is self-compatible due to a mutation in mating type genes and produces dikaryotic cells with two identical nuclei. In response to light and nutritional cues the dikaryon forms sexual fruiting bodies, in which a complex morphogenetic process culminates in meiosis and spore formation (23). While the importance of light and starvation on fruiting body development are well-known (31), the molecular underpinnings of these responses and their relationship to fruiting body development are poorly understood.

In this study we assembled a chromosome-level genome for the *C. cinerea* Amut1Bmut1 #326 strain, sequenced full-length transcripts using Oxford Nanopore and PacBio isoform sequencing, and profiled its transcriptome across a broad panel of conditions using QuantSeq. Long Nanopore and PacBio reads provide an accurate picture of full-length transcripts and their diversity, which allowed us to improve gene models, annotate uORFs, intergenic regions, 5’-, 3’ UTRs, and polyadenylation sites. Long-reads also helped to correctly identify microexons (32), which can have regulatory roles in vertebrates (33–36), insects (37, 38), and plants (39–41) but are barely studied in fungi. We amalgamate these annotations with transcriptional profiling of densely sampled time course experiments, to improve the functional annotation of *C. cinerea* genes. Using these data we show that events preceding fruiting body development of *C. cinerea* comprise a mixture of starvation and light responses as well as morphogenesis. We disentangled the gene sets belonging to each of these phenomena, and provide consensus gene lists underlying starvation, light response, and the differentiation of aerial and submerged mycelium. Finally, we made these resources openly available as a website (mushroomdb.brc.hu) for use by the community.

## Results

### New assembly for *C. cinerea*

We sequenced and assembled the 13 chromosomes of the homokaryotic Amut1Bmut1 #326 strain of *C. cinerea* genome into 23 scaffolds using PacBio reads, with an estimated 75x coverage (Figure 1/A). Based on the scaffold length distribution a nearly 10-fold decrease is observable between the fifteenth longest scaffold (1,097,509 bp) and the following scaffold (162,061 bp). After comparison of the draft genome with the available chromosomal level Okayama 7 #130 (26) and Amut1Bmut1 #326 (29) genome assemblies we found that the 15 largest scaffolds corresponded to the 13 chromosomes of *C. cinerea*, with two chromosomes each broken into two contigs. These were joined, based on a previous Nanopore-based assembly (29) that contains a linker sequence of 1000 unknown nucleotides (N1000). The size of the resulting new assembly (CopciAB V2) is slightly shorter than the previous Nanopore-based assembly; however, both of them are longer than the Okayama 7 and the published Illumina-based Amut1Bmut1 assembly (CopciAB V1) (Figure 1/B).

**Figure 1.**
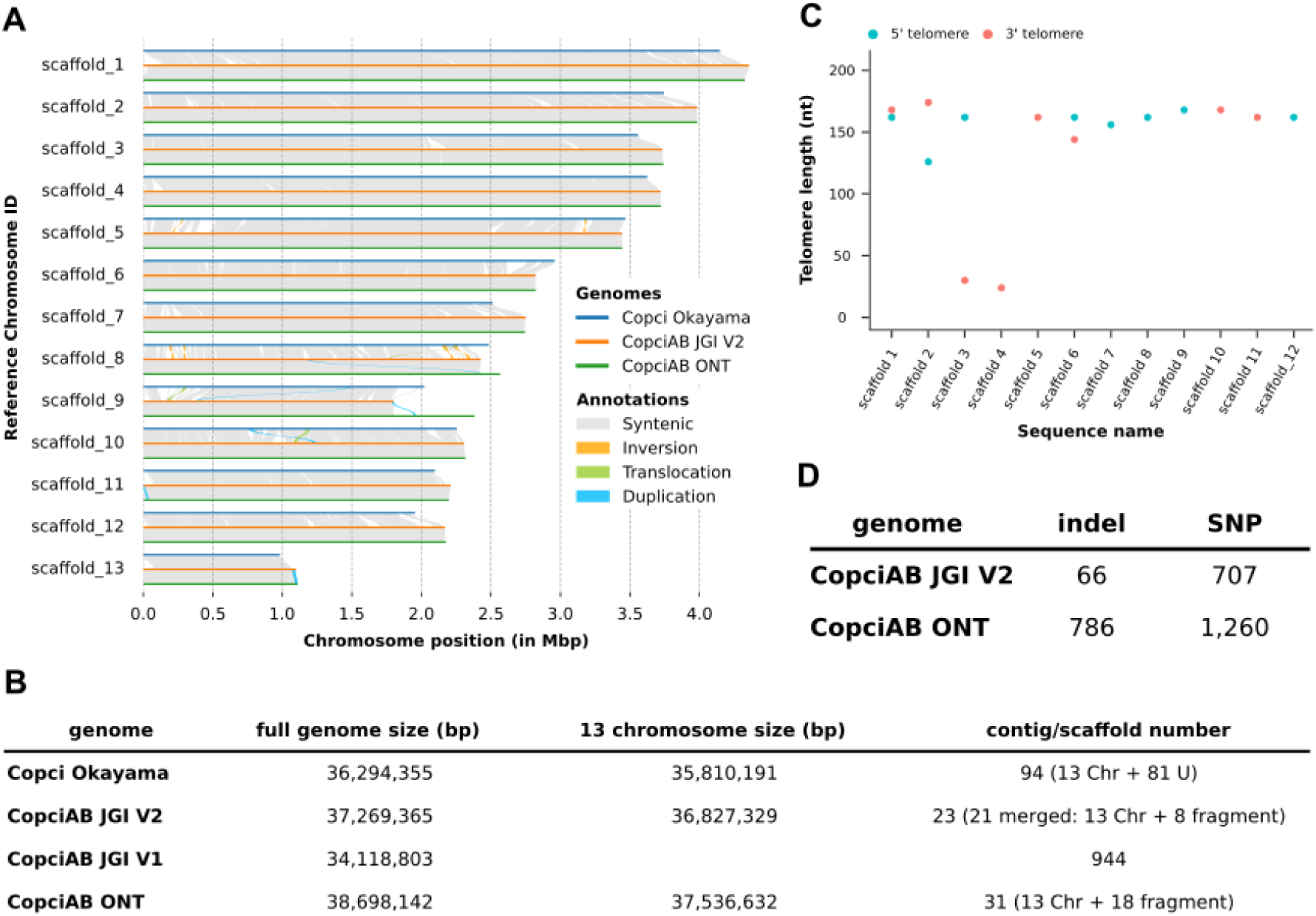
Characterisation of the new genome assembly in the light of the previous assemblies. **A.** Observed structural variance between the 13 chromosomes of the three *C. cinerea* genomes. The Nucmer function from the MUMmer v4.0 package was used for the comparisons. SyRI was used to identify genomic rearrangements and local sequence differences.Visualization was performed with plotsr. **B.** Comparison of the genome size and scaffold numbers of the available *C. cinerea* genomes. Copci Okayama stands for the short-read based *C. cinerea* Okayama assembly, CopciAB JGI V1, V2, ONT for the short-read, PacBio and ONT based assembly of the *C. cinerea* Amut1Bmut1 strains, respectively. **C.** Identified telomere repeat sequence length distribution **D.** The number of indels and SNPs occurring in the two chromosome-level CopciAB genome assemblies. Variants were called in the assemblies based on Illumina genomic read mapping.

Whole genome alignments showed a 99.11% and 99.98% average identity between CopciAB V2 and the Okayama and Nanopore-based *C. cinerea* assemblies, respectively. Analyses of structural variations between the three assemblies revealed high synteny which is more pronounced between the CopciAB V2 and the Nanopore-based genome, probably because these are from the same strain (Figure 1/A). Mapping Illumina reads on the Nanopore-based AmutBmut assemblies revealed that our new assembly contains at least tenfold fewer indels and twofold fewer SNPs than the previous Nanopore-based assembly (Figure 1/D).

In 12 of the 13 chromosomes, telomeric on at least one end repeat were identified; four chromosomes have both telomeres (Figure 1/C). On average the telomere sequences are 150 bp long (25 complete consecutive repeats) and unlike the budding and fission yeast, the tandemly repeated sequence [5’-(CCCTAA)_n_/(TTAGGG)_n_-3’] is identical to repeats from other filamentous fungi (42) and the typical human sequence (43–45). Identical telomere repeats were also identified in the case of other basidiomycetes, *Ustilago maydis* (also 150bp)*, Armillaria ostoyae, Pleurotus ostreatus* (46–48) and in other diverse fungi (49, 50).

### Genome annotation using Nanopore and PacBio cDNA-Seq

To create the most complete annotation possible, we used Nanopore and PacBio IsoSeq cDNA-seq datasets obtained from libraries enriched for full-length transcripts. We also integrated information from the two available annotations of *C. cinerea* Amut1Bmut1 #326 genomes (28, 29), followed by manually checking and correction of misidentified splice sites, transcript flanking regions and CDSs, among others. In this way, we managed to compensate most of the weaknesses of the two sequencing techniques, such as the lower read output of the PacBio IsoSeq, the lower fraction of longer transcripts in the Nanopore output (51, 52) or the truncation of transcripts with internal poly(A) or poly(T) runs by both techniques (53, 54).

For the ONT dataset, five separate samples were collected from different *C. cinerea* developmental stages, while for the PacBio Iso-Seq dataset transcript diversity was captured by pooling various tissue types. To improve ONT reads we used previously published Illumina reads (24). As a result, the quality of polished reads approaches that observed for PacBio reads (Supplementary Figure 1/B). After polishing we obtained approximately 4 million high quality reads per sample using Nanopore, in total ∼20 million reads. Over 4 million reads were gained from Iso-Seq (Supplementary Figure 1/A). The median read length in the ONT datasets is roughly half (943 bp) that of the IsoSeq dataset (1,867 bp), likely because PacBio’s IsoSeq protocol is optimized for transcripts centered around 2 kb when no size selection is used.

For gene model prediction, we combined numerous sources of information and manual curation steps (see Methods), with ONT and Iso-Seq transcripts forming the core of the annotation, supplemented with gene models from previous *C. cinerea* Amut1Bmut1 #326 annotations (28, 29). As a result, 13,617 gene models with 14,750 transcripts were predicted. Of these, 11,583 had support by at least one long-read dataset. The resulting number of genes is slightly lower than in previous annotations (28, 29), possibly because we excluded genes that had neither functional annotations, nor detectable expression, or homology outside *C. cinerea*. BUSCO indicates that the new annotation is highly complete, with minor improvements over previous *C. cinerea* genomes (Table 1). The average number and length of CDSs, exons and introns are higher than in previous annotations (Table 2). The long-read cDNA data allowed us to annotate untranslated regions (UTRs), with 12,011 and 12,130 genes having 5’- and 3’- UTRs, respectively (Table 2).

**Table 1.** BUSCO-based comparison of the three available *C. cinerea* AmutBmut genomes. Completeness of the annotation of CopciAB strains was determined using the basidiomycota_db10 database of BUSCO.

| Description | JGI V1 | JGI V2 | ONT |
| --- | --- | --- | --- |
| **Complete BUSCOs (C)** | 1,723 (97.7%) | 1,761 (99.8%) | 1,698 (96.3%) |
| **Complete and single-copy BUSCOs (S)** | 1,715 (97.2%) | 1,750 (99.2%) | 1,681 (95.3%) |
| **Complete and duplicated BUSCOs (D)** | 8 (0.5%) | 11 (0.6%) | 17 (1%) |
| **Fragmented BUSCOs (F)** | 23 (1.3%) | 1 (0.1%) | 50 (2.8%) |
| **Missing BUSCOs (M)** | 18 (1%) | 2 (0.1%) | 16 (0.9%) |

**Table 2.** Properties of *C. cinerea* protein coding gene models of CopciAB V2 compared with prior genome annotation versions. The term “genes” is defined as the complete, unmatured mRNA transcript, which consists of exons and introns. Exons describe the mature part of the mRNA, which may or may not code for a polypeptide. The CDS, or coding DNA site, is the coding part of the exon. The length is given in base pairs (bp).

|  |  | JGI V1 | JGI V2 | ONT | Okayama |
| --- | --- | --- | --- | --- | --- |
| gene | number | 14,242 | 13,617 | 15,250 | 13,393 |
|  | average length | 1748.24 | 1981.68 | 1796.85 | 1763.01 |
| cds | number | 75,485 | 75,118 | 80,939 | 75,320 |
|  | average number | 5.30 | 5.52 | 5.31 | 5.62 |
|  | average length | 1300.33 | 1351.49 | 1281.34 | 1385.96 |
| exon | number | 77,000 | 78,990 | 83,580 | 75,796 |
|  | average number | 5.41 | 5.80 | 5.48 | 5.66 |
|  | average cumulative length per gene | 1435.96 | 1667.23 | 1485.25 | 1426.82 |
| intron | number | 62,758 | 65,373 | 68,510 | 62,403 |
|  | average number | 4.41 | 4.80 | 4.49 | 4.66 |
|  | average cumulative length per gene | 312.28 | 314.44 | 311.61 | 336.19 |
| 5'UTR | number of genes with utr | 5,759 | 12,011 | 5,560 | 2,071 |
|  | percent of genes with utr | 40% | 88% | 36% | 15% |
|  | average utr length of genes with utr region | 153.76 | 147.54 | 253.60 | 89.53 |
| 3'UTR | number of genes with utr | 5,728 | 12,130 | 5,100 | 2,111 |
|  | percent of genes with utr | 40% | 89% | 33% | 16% |
|  | average utr length of genes with utr region | 182.62 | 208.36 | 329.23 | 171.39 |

### Small exons are the hallmark of complex gene models

We identified several extremely short, internally located exons, which are often referred to as microexons (32) in the literature. Correct identification of these small exons, especially those below the limit of mapping software (<15 nt), can be particularly important as their omission in the annotation can strongly influence the gene model. In the new annotation 1,165 gene models (∼9%) have at least one internal microexon of ≤15 nt; in 835 cases (∼6 %), the omission of one of these would create a premature stop codon. Notably, 89 gene models have at least one exon ≤3 nt (1 nt: 13 exons in 13 genes, 2 nt: 57 exons in 50 genes, 3 nt: 27 exons in 26 genes) and of these, an alternative stop codon is created in 62 gene models when the microexon is omitted. We found that microexons are enriched in certain gene families, among others, two families containing domains of unknown function and in cytochrome P450 encoding genes (Supplementary Figure 2) consistent with previous reports from fungi (55).

### Conserved genomic elements in the new *C. cinerea* annotation

The long-read-based gene models were used to identify conserved genomic elements located around protein coding genes. First, in ± 50 nt flanking the Transcription Start Site (TSS) of long-read based gene models with 5’UTR region (n:11,483) we identified patterns consistent with the eukaryotic Initiator sequence (Inr) minimal consensus (C/T)(<u>A/G</u>), where <u>A/G</u> is the TSS (Figure 2/A), in line with previous results in other organisms (56–60). Within 50 nt upstream of the TSS, 20% (n:2,323) of genes have at least one TATA box-like sequence (TATANN), 1,376 of which had a canonical TATA box (TATATA or, TATAAA). Of the identified canonical TATA boxes, 83% (n: 1,151), start between −38 and −31 nt (most frequent: −34 nt). Noncanonical TATA boxes were also enriched in these locations. The location of TATA boxes is similar to that observed in *Aspergillus nidulans* (59) and *Schizosaccharomyces pombe* (60) where TATA boxes are located −45 to −30 nt, or −32 to −25 nt upstream of the TSS, respectively.

**Figure 2.**
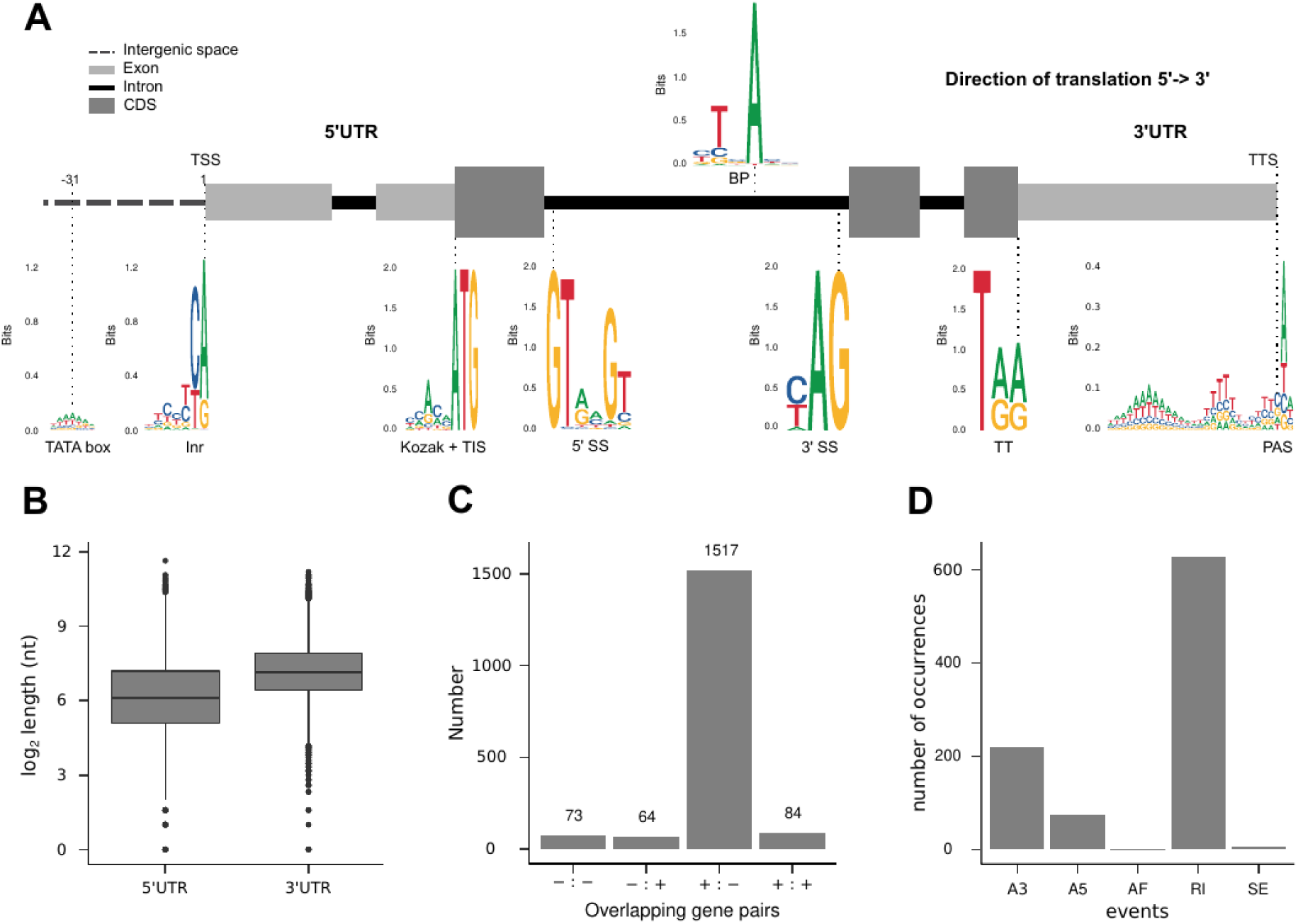
Presentation of the characteristic features of the predicted gene models of *C. cinerea*. A. Schematic representation of a typical *C. cinerea* protein coding gene model, highlighting conserved motifs. TATA box and initiator (Inr) consensus motifs typically associated with highly expressed promoters and located in close proximity to transcription start site (TSS). Kozak sequence context localizes adjacent to the translation initiation site (TIS) and regulates its selection. Conserved 5’ splice site (5’ SS), branch point (BP) and 3’ splice site (3’SS) motifs coordinate the spliceosome activity. Translation termination (TT) marks the sequence context defined by the stop codons. The transcription termination site (TTS) marks the 3’ end of the transcript, which is defined by the polyadenylation site (PAS). B. UTR length distributions plotted as boxplots. The bottom of the box represents the first quartile of the data, the black line in the middle of the box represents the median of the data, and the top of the box represents the third quartile of the data. C. The distribution of the overlapping gene pairs in four possible combinations of gene orientation. The combinations were indicated in the following way: −/− expressed from the antisense strands in the same direction, +/+ expressed from the sense strands in the same direction, −/+ expressed divergently from antisense and sense strands, and +/- expressed convergently from the sense and antisense strands. D. Distribution of the alternative splicing events across alternative 3’ splice site (A3), alternative 5’ splice site (A5), alternative first exon (AF), retained intron (RI) and skipping exon (SE).

We also identified signals for the Kozak consensus sequence in the vicinity of the most likely AUG start codon in the top 10% of all transcribed genes based on long-read support (n: 1,096) (Figure 2/A), which is similar to that of other fungal species (56). Regarding splice sites, 99.8% found in 10,890 gene models with at least one splice site belonged to the GT-AG, GC-AG and AT-AC subtypes (Figure 2/A), which is consistent with the literature (61). The translation termination site is represented by one of the three stop codons (UAA, UGA, UAG), and for the long-read-based annotations these codons were almost evenly used (30.4%, 37.8%, 31.8%, respectively) (Figure 2/A). We detected motifs around the Transcription Termination Site (TTS) (Figure 2/A) similar to those observed in other fungal species especially in *Schizosaccharomyces pombe* (62, 63), *Magnaporthe oryzae* (64), *Fusarium graminearum* (16), *Saccharomyces cerevisiae* (63), as well as in other multicellular organisms (65).

#### uORFs

Reliable identification of 5’UTRs provided an opportunity to investigate the occurrence of uORFs in *C. cinerea*. We considered all alternative ORFs whose start codon is within the 5’UTR region as uORFs as long as they encode at least one amino acid before a stop codon (i.e., at least 9 nt long) (66). We identified a total of 8,704 uORFs, with median predicted peptide length of 19 aa (mean 29.22 aa), found in 26% (n: 2,982) of the 11,483 predicted genes. The largest proportion (46%) of these genes have a single uORF. The observed proportion of genes possessing uORFs (56, 66, 67) and the median peptide length of the uORFs is not unusual compared to previously described uORFs (66). We identified a putative homolog of the arginine attenuator peptide (AAP), which was first described in the 5’UTR of the arg-2 gene of *Neurospora crassa* (68, 69), in the 5’UTR of the orthologous *C. cinerea* gene (CopciAB_446268) (Supplementary Figure 3). These data indicate that, similarly to other fungal and eukaryotic genomes, uORFs are widespread in 5’UTRs of *C. cinerea* genes and that some uORFs are highly conserved across fungi.

#### UTRs

Across all gene models (n=13,617), 88% and 89% of the genes have a 5’UTR and 3’UTR annotation, respectively. The median length of the 3’UTR regions is at least twice larger than that of 5’UTRs (5’UTR: 68 nt, 3’UTR: 141 nt) (Figure 2/B). We found 1,738 long-read based adjacent gene pairs with overlapping UTR annotations, with a median overlap of 99 nt. 1,517 of these overlaps were found between convergently oriented gene pairs (+><-), whereas in other orientations the overlap was considerably less frequent (Figure 2/C). Such overlapping UTRs in convergently oriented gene pairs are a common feature in fungal genomes and can affect their transcription and translation (13, 14, 70, 71).

#### Transcript isoforms and alternative splicing

As described earlier, an appropriately sensitive method will eventually detect splice variants for every multi-exon gene (72), not all of which may be biologically relevant. Therefore, only gene isoforms with a read support of at least 10% of total read for each gene were considered here. Using this conservative approach, 1,053 genes, 7.7% of all genes, were found to have at least one isoform, which is consistent with previous studies (73), but considerably fewer than Illumina-based splicing inferences from two recent studies (24, 74). The relative frequencies of splicing events was consistent with those reported earlier (16, 73, 75), with retained introns being most common (67.5%), followed by alternative 3’ splice site (23.5%), alternative 5’ splice site (8.2%), exon skipping (0.5%) and alternative first exon (0.2%) (Figure 2/D).

### Alternative polyadenylation

To annotate and explore the use of polyadenylation sites (PAS) in *C. cinerea*, we profiled gene expression across 67 different conditions covering several aspects of the biology of the fungus. We used QuantSeq, a 3’ sequencing approach that yields information on the junction of 3’UTRs and poly(A) tail (76) and thus can provide PAS information (56). In the first round all the available QuantSeq reads were used to identify 339,309 PAS. Because we applied stringent filtering criteria (see Methods), we could use only 356 million of the 27 billion QuantSeq reads. Nearby PAS were clustered into single polyadenylation site clusters (PACs) by allowing a maximum of 10 nt intra-cluster distance, yielding 76,879 PACs. Within a PAC the PAS with the highest read support serves as the representative PAS for the cluster (Figure 3/A), and we used its genomic coordinate for defining a specific PAC.

**Figure 3.**
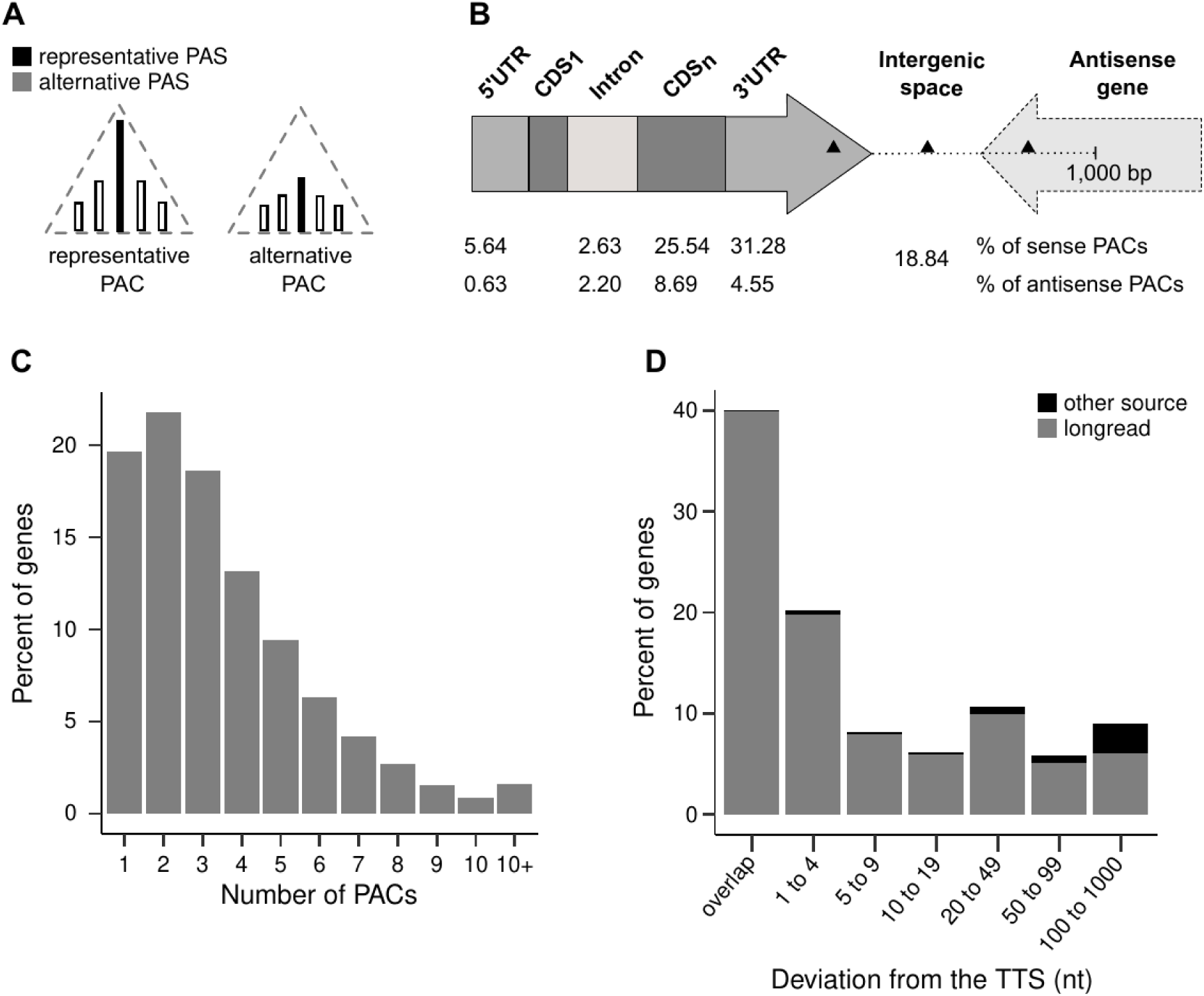
Description of the relationship between the predicted gene models and the PACs. A. Schematic diagram illustrating the hierarchy of PASs within PACs. The representative PAS with the largest read support is colored black. The best supported PAC in the case of APA is marked by gray background. B. Proportions of 76,879 PAC mapping to parts of a gene model and intergenic region. 40,994 PAC, marked with black triangle, were assigned to the genes when located in the 3’UTR of the gene, in the intergenic region or in the genic region of downstream antisense genes <1,000 bp from the gene’s TTS. Note that, if the gene is followed by another in the same orientations, then different rules were used (see Methods). C. Proportion of genes with different number of PACs. D. Percentage of genes whose TTS is within a certain distance from the PAC site in genes of different origins in the annotation

PACs are enriched in the “sense” orientation in 3’UTRs (31.28%) and CDS (25.54%) regions of the gene models (Figure 3/B). PACs annotated to non-3’UTR parts of the gene models may be results of internal priming or template switching at sites with >3-5 adenines (54), to which the QuantSeq protocol is susceptible, and can thus produce false-positive PAC predictions. The presence of spurious PACs is suggested by the ∼3x higher median expression of the representative PAS site of PACs in the 3’UTRs than that in the other annotation entries (Supplementary Table 1). Therefore, we further filtered PACs based on localization, orientation, keeping in mind that due to the frequent overlap of 3’ UTR regions between fungal genes, relevant PACs can also be located in neighboring genes. In this way, we obtained 40,994 PACs which were assigned to 11,628 gene models. Over 80% of the genes with an assigned PAC have more than one PAC and we thus considered them as genes with putative alternative polyadenylation (APA) (Figure 3/C). For each APA gene, the PAC whose representative PAS had the highest count is hereafter referred to as the best-supported PAC (Figure 3/A).

Transcription termination site coordinates of genes from the long-read-based annotation perfectly overlap with the coordinates of the best-supported PAC in ∼40% of genes and deviates from it by at most 4 nt in another 19.75% (Figure 3/D). In contrast, the distances of non-long-read-based gene models from their best-supported PAC are considerably larger (Figure 3/C), perhaps due to the possible absence or misannotation of the 3’ UTR regions.

To understand the diversity of PAC usage in APA genes across biological conditions, we calculated the evenness in each experiment for each APA gene with an expression ≥10 count per million (CPM). Figure 4/A-C shows that there is a significant negative correlation between evenness and gene expression level, that is, the higher the gene’s expression, the less diverse its PAC usage is. This could indicate that for each gene a certain PAC is preferred over others, which is especially notable for highly expressed genes. To gain insight into the preferred PAC among the different biological conditions, we ranked PACs based on their relative expression, calculated as the expression value of the PAC divided by the summed expression of all PACs of the gene (77).

**Figure 4.**
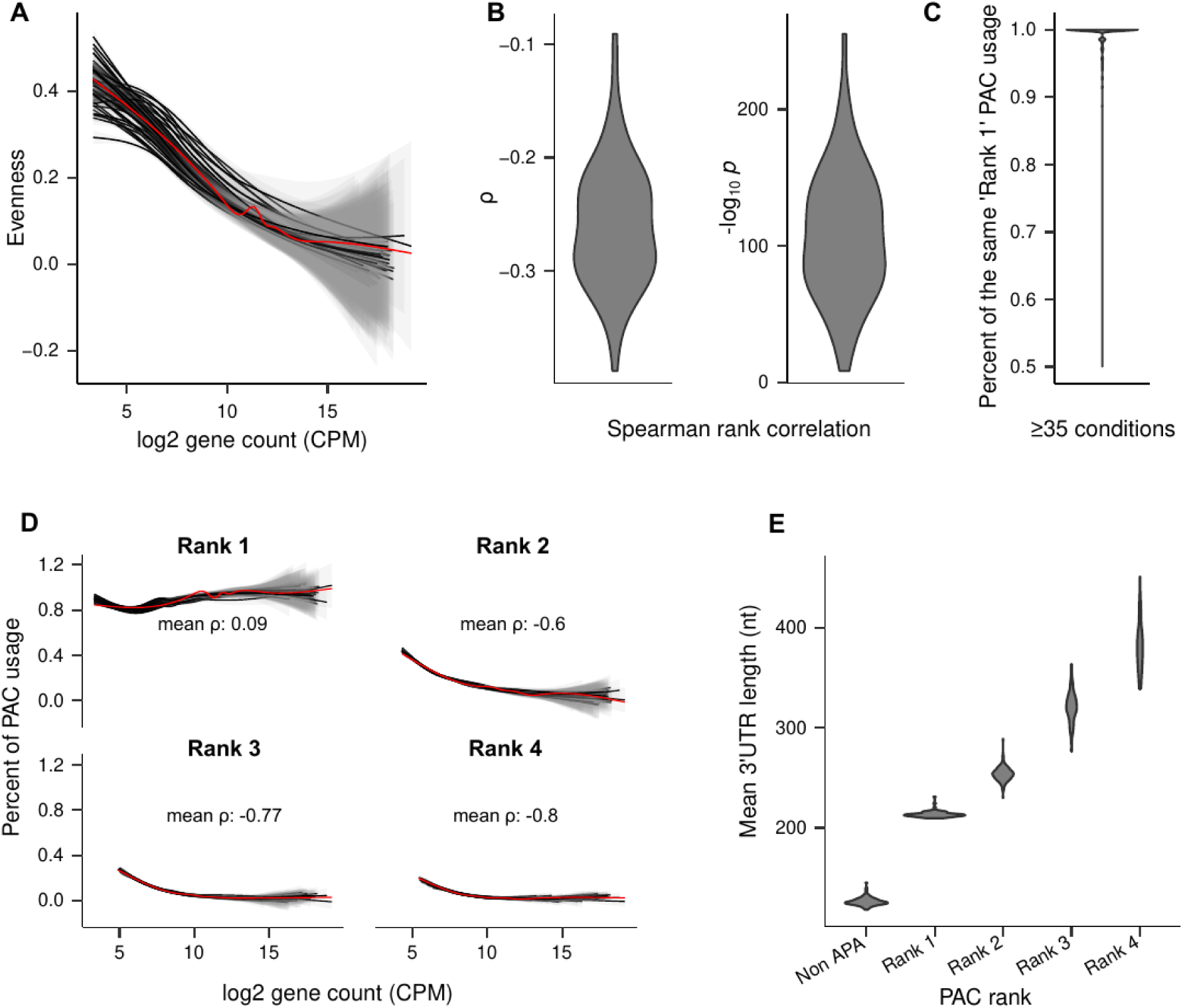
Characterization of PACs use preference in *C. cinerea*. A. Negative correlation between evenness of PAC site usage and gene expression in APA genes. Note that the higher the expression the less diverse PAC usage is. Lines represent 67 biological conditions analyzed by QuantSeq. The red line represents a mean across all biological conditions. B. Violin diagrams of the distribution of the Spearman rank correlation rho (ρ) and the − log_10_ *p* values of the correlation between evenness and log_2_ gene count across the 67 biological conditions. C. Violin diagram showing the relative usage of the PAC with highest read support (Rank 1) for APA genes expressed in at least 35 biological conditions. Only APA genes with at least 1 PAC having >10 CPM were considered. D. The correlation between the frequency of a given PAC rank and gene expression. The black lines represent 67 biological conditions. The red line represents a mean across all biological conditions. E. The association between PAC usage and mean 3’ UTR length across non-APA and APA genes. Note the clear correlation between rank (i.e., decreasing expression) and 3’ UTR length. Violin plots represent 3’ UTR length distributions across biological conditions.

We found that for most genes the transcriptional machinery used the same, most highly expressed (Rank 1) PAC under all biological conditions we tested (Figure 4/C). We also found that this correlation held strongly for highly expressed genes (Figure 4/D).

When we checked the mean 3’ UTR length variation in the first four APA ranks together with that of non-APA genes (≥10 CPM), we found that the non-APA genes had shorter 3’ UTRs region followed by the highest expressed PAC (Rank 1) and other PACs in order of decreasing read support (Rank 2 to Rank 4) (Figure 4/E). Thus, there is a clear correlation between PAC use frequency and distance from the CDS; that is, the higher the relative expression of the PAC, the shorter the resulting 3’ UTR. This suggests that *C. cinerea* most often expresses the transcript with the shortest 3’ UTR version. At the same time, we did not find significant variation of PAC usage across biological conditions, which suggests that APA is a prevalent though probably not influential phenomenon in *C. cinerea*. This is consistent with views that suggest that APA represents “transcriptional noise” and not an actively regulated process (77). Collectively, the annotated polyadenylation clusters correspond very closely with the predicted TTS of the gene models derived from long reads, indicating that our long-read-based gene models, including their 3’ UTRs represent robust annotations. In terms of alternative polyadenylation, it appears that most genes, especially highly expressed ones, prefer one PAC, and mainly the one that results in the shortest 3’ UTR.

### Expression data disentangle light and starvation responses and morphogenesis

In contrast to Ascomycota, where extrapolations from model organisms and a long tradition of both forward and reverse genetics studies yielded functional information for a considerable proportion of genes, Basidiomycota gene function is notoriously poorly understood. To mitigate this, we took advantage of the improved annotation of *C. cinerea* and profiled gene expression under a broad panel of conditions (201 samples for 67 conditions), covering basidiospore and oidium germination, mycelium growth in the dark and after light induction, hyphal knot, sclerotium and fruiting body development, carbon starvation, as well as 11 different stress conditions. These data were used to generate functional insights into the *C. cinerea* gene repertoire. Differential gene expression analyses identified a total of 12,569 differentially expressed genes (DEGs, BH adjusted p ≤0.05, fold-change ≥2) (Supplementary Figure 4), indicating that the 67 conditions induced significantly altered expression of 92.3% of all genes. Here we focus more closely on densely sampled time course data for light induction, morphogenesis and starvation (Figure 5/A). Fruiting body development in the basidiomycetes happens in response to changing environmental variables (31, 78), of which the most important are carbon starvation and light (31). However, under standard fruiting conditions distinguishing the specific gene expression responses attributable to starvation and light induction from those associated with morphogenesis has not been possible. To this end, we obtained consensus gene lists by comparing differentially expressed genes in two starvation-inducing experiments, after light-induction and two mycelial types (aerial and attached mycelium). Figure 5/B shows how comparisons of DEGs from these experiments yielded functional gene groups we designate as ‘core starvation response’, ‘core light response’ and ‘core mycelium differentiation genes’ in.*C. cinerea*.

**Figure 5.**
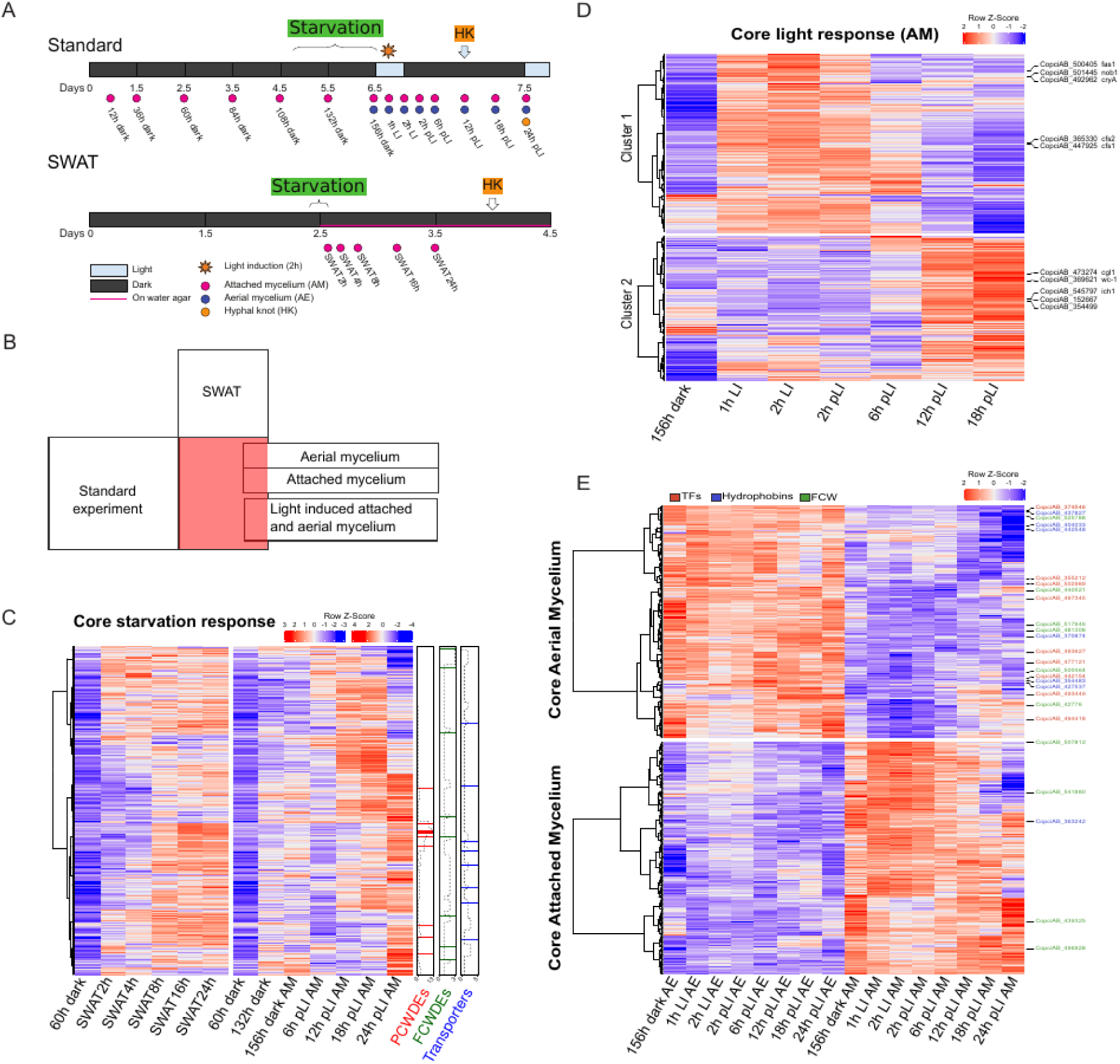
Illustration of the experimental design used to define core gene sets whose expression profile is shown in the form of a heatmap. A. Graphical representation of the experimental conditions used for light induction, starvation and mycelial type. The day-light cycles are marked on the strip with dark-light segments. The reference times are marked below the strip. The collected samples are shown with a colored dot below the time references, where the colors represent the tissue type of the collected sample. B. The stylized Venn diagram illustrates the overlaps between the sample sets which were made. The red square marks the core starvation response, which is present to some extent in all sample groups. C. Heatmap representation of the log_2_CPM expression values of the core starvation response genes of the water agar transfer (SWAT, left) and half-glucose YMG medium-induced starvation (standard, right) experiments. The columns on the right mark the position of the plant cell wall degrading enzyme genes (PCWDEs, red), fungal cell wall related enzymes (FCWDEs, green) and transporter-encoding genes on the heatmap. To visualize the distribution of the genes, a rolling window with a width of 20 genes and a length of 20 steps was used and shown with a dashed line. DEFINE all abbreviations here D. Heatmap representation of the log_2_CPM expression values of the core light response genes in the AM samples. Genes were hierarchically clustered into two groups using the WardD method based on the Euclidean distance between the Z-score-normalized log_2_CPM gene expression values. Genes identified in the literature as early and late light-induced genes are marked on the right side of the figure. E. Heatmap representation of the log_2_ CPM expression values of the core aerial and attached mycelium specific genes in the aerial and attached mycelium samples. Putative TF-, hydrophobin- and FCWDE-encoding genes are marked with red, blue and green colors on the right side of the figure.

#### Carbon starvation

We induced starvation by growing *C. cinerea* on Yeast-Malt-Glucose (YMG) plates with reduced (0.2%) glucose content (called “Standard” experiment) (28) and by transferring 60 h old dark-grown colonies to water-agar in the darkness, which contained no nutrients (hereafter called “SWAT” experiment) (Figure 5/A). In both the Standard and SWAT experiments, DEG analyses were run with the 60 h dark mycelium sample as the reference. In the standard experiment nutrients deplete gradually, whereas in the SWAT experiments nutrient depletion is abrupt, expected to induce a stronger starvation response. Accordingly, in SWAT experiments most gene expression changes are observable at 8 h, whereas in the standard experiment upregulation patterns are more diverse, which could stem from a combination of milder starvation, light irradiation and tissue differentiation. We found 3,242 and 2,348 significantly upregulated genes (*p* ≤0.05, fold change ≥2) in the Standard and SWAT experiments, respectively. Of these, 1,759 genes were upregulated in both experiments and are hereafter refer to as “core starvation response genes” (Supplementary Table 2).

GO enrichment analyses provided a clear signal for the upregulation of genes encoding plant cell wall degrading enzymes (PCWDEs) and transporters, among others (Supplementary Figure 5). As expected, several enriched GO terms related to carbohydrate utilization were linked to PCWDEs. Based on these findings and previous considerations of starvation, we prioritized plant cell wall degrading CAZymes (PCWDEs) and transporters, as well as autophagy and fungal cell wall degrading CAZymes (FCWDEs) for deeper scrutiny.

The two most abundant functional groups in the core starvation response, consistent with GO enrichment results, were PCWDEs and plasma membrane transporters. Out of the 213 PCWDEs and 375 transporters of *C. cinerea*, 136 and 134 were upregulated in at least one of the starvation experiments and 96 and 55 (Supplementary Figure 6) were among the core starvation response genes, respectively. We also compared these gene lists with *C. cinerea* genes upregulated in carbon-catabolite repressor (Cre1, CopciAB_466792) gene deletion strains (79) and found that starvation-induced PCWDEs and transporters overlap considerably with those upregulated in the *Δcre1* strains. Of the 62 PCWDEs and 19 transporters upregulated in *Δcre1*, 52 (83.9%) and 10 (52.6%) were also part of the core starvation response, respectively.

Core starvation-responsive PCWDEs had diverse predicted substrates and included 67 cellulose (e.g. AA9, CBM1, GH3, GH5, GH6, GH7, expansins), 30 hemicellulose (e.g. GH11, GH31, GH93), 16 pectin (GH28, GH88, various PL families) but none of the few lignin degradation related genes of *C. cinerea*. Of the 364 transcription factors (TFs) identified in *C. cinerea*, the ortholog (CopciAB_447627) of the cellulase-activating *roc1* transcription factor of *Schizophyllum commune* (80) was part of the core starvation response genes together with additional 39 TFs. Five of these TFs genes, including *roc1*, were also upregulated in the ΔCre1 strains indicating that regulators of PCWDEs also respond to starvation.

Among transporters, the core starvation response contained many members of the Major facilitator superfamily (MFS) (IPR036259) some of which are putative orthologs of experimentally characterized Ascomycota glucose and xylose transporters, including HGT1/HGT-1 of *Candida albicans* and *N. crassa* (CopciAB_501903) or XtrD of *Aspergillus nidulans* (CopciAB_358584). MFS transporters in the core starvation response and upregulated in the ΔCre1 strain included putative orthologs of transporters involved in the transport of cellulose degradation intermediates. For example, CopciAB_496081 and CopciAB_496145 encode putative orthologs of the *N. crassa* cellodextrin transporter CLP1 and hexose transporter CDT-2 (81, 82) (Supplementary Figure 7).

The core starvation response contains several genes which were considered essential for fruiting body formation. The *dst-2* gene (CopciAB_361584) (83) with a FAD domain was hypothesized to be a photoreceptor for blue light and has been proven essential for photomorphogenesis in *C. cinerea*. Its upregulation in the SWAT experiments is surprising given that these experiments did not receive light. Two fungal galectins, Cgl1 and Cgl2 (CopciAB_473274, CopciAB_488611) (84), which are considered to be involved in hyphal knot induction were responsive to starvation. This is in line with publications (84, 85) showing the link between starvation and fruiting body induction. None of these genes was upregulated in the ΔCre1 strain.

One of the hallmarks of starvation is autophagy and hyphal autolysis, which may be associated with increased expression of autophagy and fungal cell wall remodeling CAZyme genes (86–88), respectively. In *C. cinerea* we found 12 putative orthologs (reciprocal best hits) of 35 previously reported autophagy genes of *S. cerevisiae* (88). Eight of these showed increasing expression 2h after transfer to water-agar (Supplementary Figure 8), however, only one of them (CopciAB_375414) was significantly differentially expressed. Of the 138 *C. cinerea* genes encoding CAZymes with an activity on fungal cell wall, 52 and 59 were upregulated in the SWAT and standard experiments, respectively, and 40 were part of the core response (Supplementary Figure 6). This suggests that there is active remodeling and/or digestion of the fungal cell wall, which could indicate autolytic processes in starving *C. cinerea* cultures.

Taken together, we defined a core set of genes induced upon starvation, which can be used to understand how the fungus prepares for fruiting body development, the main inducer of which is nutrient depletion. These genes indicated that upon starvation, *C. cinerea* was probably released from carbon catabolite repression, transcribed genes associated with plant cell wall degradation and morphogenesis and showed signs of autophagy or autolysis. These observations are similar to those made in starving Ascomycota cultures (86, 88, 89). Autophagy has also been detected upon prolonged starvation in the basidiomycete *Paxillus involutus* (87). In the ascomycetes the carbon starvation response overlaps with the response to lignocellulosic carbon sources, in that it comprises the induction of several PCWDEs (86, 90, 91). We found similar patterns in *C. cinerea* and hypothesize that this is because, upon glucose depletion, the fungus explores available alternative carbon sources by expressing a broad panel of hydrolytic enzymes. Such scouting enzymes may generate simple sugars that transporters import into the cell, which may trigger positive regulatory feedback loops that strengthen the transcription of genes related to the carbon source found (89, 92).

#### Light-induced genes

We examined the effects of light induction by using the synchronized growth protocol developed previously (28), which involved a 2 h light induction followed by 24 h in the dark and then alternating 12 h light/dark cycles. We sampled aerial and attached mycelium separately for eight time points (Figure 5/A), and identified light-induced genes by comparing all time points to 6.5-day-old dark-grown aerial or attached mycelium as reference (called 156 h dark, see Figure 5/A). We identified 2,453 and 2,065 DEGs in the attached and aerial mycelium samples (Supplementary Table 3), respectively. Aerial and attached mycelia shared 1,251 DEGs, indicating high similarity in light response irrespective of mycelial type. We used the union of 2,453 and 2,065 genes and removed genes that belong to the core starvation response to identify 2,340 genes specifically induced by light irrespective of mycelial type, but not by starvation. We hereafter refer to these as “core light induced genes” (Figure 5/B, Supplementary Table 3).

To understand the tempo and mode of light induced gene expression changes we analyzed the expression of 2,340 core light induced genes in attached mycelium samples (Figure 5/D), and noted that aerial mycelia show similar, though noisier patterns (see Supplementary Figure 9). An expression heatmap revealed two clear waves of gene upregulation, one evident 1 h after the start of light induction and decreasing expression after 6 h post light induction (pLi) and another wave starting around 12 h pLi (Figure 5/D). Hierarchical clustering recovered two clusters corresponding to these two waves. We hereafter refer to cluster 1 (1,291 genes) and cluster 2 (1,049 genes) as early and late genes, respectively, which probably correspond to the two waves of light responsive gene induction reported earlier (85). The expression of early genes started to decline relatively quickly, consistent with photoadaptation mechanisms that inhibit White Collar Complex (WCC) activity. We note that the first hyphal knots emerged at 12 h pLI, so it is possible that late genes include ones related to the onset of hyphal knot development.

GO analyses revealed a significant effect of early light induction on peptide metabolism (GO:0006518), translation (GO:0006412), ribosome biogenesis (GO:0015934), lipid metabolism (GO:0006629), DNA repair (GO:0006281), chromatin remodeling (GO:0006338), mitotic cell cycle (GO:1903047), GPI-anchor biosynthesis (GO:0006506) and mitochondrion organization (GO:0007005), among others (Supplementary Figure 10). Of these the significance of chromatin remodeling has been shown for sexual development of *C. cinerea* during the examination of the Cc.snf5 (CopciAB_365798) (93). In contrast, late light induced genes show an enrichment of different GO terms, including those related to transcription regulation (GO:0006355), fungal-type cell wall (GO:0009277), DNA packaging (GO:0044815) or G-protein signaling (GO:0007186) (Supplementary Figure 10), consistent with the appearance of the first hyphal knots 12 h pLi.

Among the early light-induced genes we found four *C. cinerea* genes (*fas1, nod1, cfs1* and *cfs2*) that were reported earlier to respond quickly to blue light (85, 94). The ortholog of the *A. nidulans* blue light sensor/photolyase *cryA* (CopciAB_492962), is part of the early gene set, while the other well characterized blue-light sensor gene, wc-1 (*dst1*, CopciAB_369621) is part of the late gene set. Similarly, *cgl2* (CopciAB_488611), which was proposed to be a marker gene of primary hyphal knots (23), was among the early light-induced genes, with an upregulation 1 h after the start of light irradiation, whereas *cgl1* (CopciAB_473274), a proposed marker of secondary hyphal knots only followed later, 12 h pLi. The wc-2 ortholog (CopciAB_8610) (95) was not upregulated in the tested samples.

We also identified diverse genes related to DNA-repair (GO:0006281) (Supplementary Figure 11), possibly for alleviating the effects of DNA damage caused by the ultraviolet component of white light. For example, we detected an ortholog of the *S. pombe* UV-endonuclease, uvdE (CopciAB_359391) and a cryA ortholog photolyase (CopciAB_492962), which is WCC-activated in *Ustilago* (96), *Schizophyllum* (97) and *Neurospora* (98) and provide protection from phototoxicity. These observations suggest that these and other early genes might be WCC-activated in *C. cinerea,* and may be part of a conserved light-responsive DNA repair network. We hypothesize that the broad upregulation of DNA repair genes is related to repairing DNA damage caused by UV or ROS generated by photons (99).

Early light-induction resulted in the upregulation of diverse lipid metabolism and genes related to Glycosylphosphatidylinisotol-anchor synthesis. Of the nine predicted fatty acid desaturases, which include linoleic acid synthases, five can be found in the early light response gene set (cluster 1) and only one among late light response genes (cluster 2). The upregulation of putatively linoleic acid producing fatty acid desaturase genes suggest that membrane fluidity is regulated in response to light, similar to what was found in fruiting bodies (78).

In summary, our data show that light induces broad transcriptional changes in *C. cinerea*, with the early light-response comprising 1,291 genes, ∼10% of all genes of the fungus. These patterns are similar to the situation reported in *N. crassa*, where an early wave of light induction is regulated by the WCC. We note that the early wave of gene expression in *C. cinerea* (0-6h pLi) is considerably slower than that of *N. crassa* (45min) (100) and more research is needed to clarify if they represent homologous responses. We hypothesize that *C. cinerea* early genes are also regulated by the WCC, even though motif analyses did not recover a motif similar to that reported in *N. crassa* (101) or *Lentinula edodes* (102). It is also noteworthy that late light induced genes show up in a consensus catalog of conserved fruiting-related genes (78), indicating that illumination induces light-responsive genes also during fruiting body development. Finally, it should be noted that here we specifically analyzed upregulated genes, analyses of light-repressed genes might yield additional insights into the photobiology of *C. cinerea*.

#### Genes related to mycelial type

In fungal colonies hyphae might be submerged in or attached to the substrate or be erect, forming the aerial mycelium. Fruiting bodies develop from hyphal knots, which in turn emerge by the branching of erect hyphae (23, 103, 104), indicating that aerial mycelium may have an important role in fruiting body formation. The transition from attached to aerial mycelium is thus an important step, however, in contrast to fruiting body development, very few reports have scrutinized it (103–105). Therefore, we analyzed gene expression separately for the aerial vs. attached mycelia at each time point in our light induction time course experiment (Figure 5/A). Since each pair of samples compared received the same light treatment, we expect DEGs to reflect differences mainly related to tissue type. We identified 501 - 1,275 and 574 - 1,135 DEGs upregulated genes in attached and aerial mycelium across the time course, respectively (Supplementary table 4).

GO enrichment analysis revealed that diverse metabolic processes dominated DEGs upregulated in attached mycelia, whereas in aerial-mycelium, terms related to transcription regulation (GO:000635), the cell wall (GO:0009277), transporters (GO:0055085) were consistently enriched, among others (Supplementary Figure 12-13). This suggests that the two mycelial types perform divergent functions; in the attached mycelium more housekeeping and metabolic functions might be active, while the transcriptome of aerial mycelium reflects differentiation and consequences of an oxygen-rich environment (e.g., response to oxidative stress). The term “Fungal-type cell wall” (GO:0009277), which is linked, among others to hydrophobin (IPR001338) upregulation was enriched in both mycelial types, however, considerably more strongly in aerial mycelium (Supplementary Figure 12-13). Aerial mycelia further showed GO enrichment signal for lipid (GO:0006629), steroid (GO:0006694) and sphingolipid (GO:0006665) metabolism (Supplementary Figure 12-13). To follow up the GO results, we analyzed the expression of genes encoding hydrophobins, cell wall synthesis and remodeling enzymes and transcription factors. Aerial mycelia displayed a considerably stronger hydrophobin gene expression than attached mycelium (Supplementary Figure 14) in line with previous studies (103, 106, 107). Hydrophobin and FCWDE-related CAZyme gene upregulation in the aerial mycelium became stronger as time progressed, possibly indicating increasing specialization at the cell surface. Generally, more TFs were upregulated in the aerial than in the attached mycelium, though the numbers converged as time progressed and at 24 h pLi the attached mycelium had more upregulated TFs than the aerial mycelium.

We next attempted to identify a core set of genes that are consistently upregulated in attached or aerial mycelium. This yielded 407 and 387 upregulated in at least five out of eight comparisons in the aerial and attached mycelia, respectively. After removing genes that are also members of the core starvation response, we obtained 197 and 247 genes (Figure 5/E, Supplementary Table 5, Supplementary Figure 15). This low overlap suggests that there is a considerable turnover of gene expression through the time points. Among these, aerial mycelia had nine TFs, six hydrophobins and six FCW genes upregulated, whereas attached mycelia had no, one, or four upregulated genes in these categories, respectively. The nine TFs consistently upregulated in aerial mycelium suggest there are specific regulatory patterns in this tissue type. One of these is orthologous to *hom1* of *S. commune* (CopciAB_493627), which was reported to regulate vegetative growth and fruiting body development (108, 109).

### Website

The *C. cinerea* 2.0 genome has been deposited in MycoCosm (3) and, to make the genome sequence, gene models, functional annotations and gene expression data accessible to the community, in a new a website (mushroomdb.brc.hu) that contains all of the above data in an easy-to-use format was generated. The website offers the possibility to identify the desired gene based on sequence similarity and keywords from the functional annotation, incorporates a genome viewer and browser (long-read alignments, Illumina and QuantSeq read coverage), data on gene conservation, orthology in other mushroom-forming fungi and tools to analyze gene expression levels in new and published datasets (24, 110–112). We anticipate MushroomDB will be a useful resource for mushroom science.

## Discussion

In this study, we produced a high-quality annotation of protein-coding genes of *C. cinerea* Amut1Bmut1 #326 by combining a new, chromosome-level assembly, Nanopore and PacBio IsoSeq cDNA-seq data, information from two available annotations and several rounds of manual correction. To improve functional descriptions of genes, their expression profile was determined under several conditions, which we used to glean novel insights into the transcriptome states related to starvation, light induction and mycelial differentiation.

The new *C. cinerea* genome eliminates several problems associated with previous annotated genomes of this species and Agaricomycetes in general. First, most Agaricomycete genomes are drafts and only a few are chromosome level assemblies (5–11), which limits their usefulness for downstream analyses. In addition, gene annotation is usually based on short-read assisted automatic pipelines, which fail to capture key gene features (e.g., APA, uORFs, UTRs) and often yield imprecise gene models; and thus incorrect or incomplete protein predictions due to challenges posed by overlapping gene models, polycistronic transcription (15, 16), or microexons, among others. It has been shown that such annotation errors can propagate across genomes (113) and can reduce the accuracy of functional annotation terms which leads to poor predictions of the biological function of genes and encoded proteins.

The new *C. cinerea* genome has better quality metrics than previous ones, such as fewer indels and mismatches, at least one telomeric repeat region at the termini of twelve of the 13 chromosomes, higher annotation completeness as suggested by BUSCO, 85% of the 13,617 gene models confirmed with at least one long read, or 5’ and 3’ UTR annotations for 88% and 89% of the gene models, respectively. Comprehensive UTR annotations enabled the detection of (un)structured elements, such as uORFs associated with 2,982 genes. By uncovering these genomic features in *C. cinerea,* this study adds the first multicellular Basidiomycota to the considerably denser array of Ascomycota (e.g. *S. cerevisiae, N. crassa, S. pombe, C. albicans*) in which these features have been already analyzed in detail, enabling comparative analyses.

For example, we confirmed the presence of an uORF in the 5’ UTR of the *arg2* gene of *C. cinerea* (69), which, together with its ortholog in *N. crassa,* indicate 500 million years of conservation for this uORF. The improved annotation enabled the detection of conserved sequence motifs (e.g., Inr, TATA-box) and will help defining cis-regulatory elements. We found a large number of microexons which, in our experience, are often overlooked by automatic annotators and can lead to premature stop codons. To further validate gene models we utilized QuantSeq’s ability to capture the junction of the 3’ UTRs and the poly(A) tail. This way we were able to confirm the PAS of the gene models and also obtained information on alternative polyadenylation (APA).

By building on the improved gene models and gene expression profiling under 67 conditions we enriched the annotation with expression-derived descriptions of gene function. Gene expression data covered light-induced developmental processes, carbon starvation as well as eleven stress conditions. We focused on a subset of these to define functional gene sets representing the core response to starvation (1,759 genes), light induction (2,340 genes) and hyphal differentiation (197 and 247 genes in aerial and attached mycelium). These three factors are usually simultaneously present to varying extents in fruiting experiments (31) and previous studies could not tease apart the transcriptomic responses attributable to them from those of fruiting body morphogenesis. The gene sets we defined displayed characteristic functional enrichment patterns and were free of contaminating effects across these three phenomena. Thus, we consider the establishment of the four core genes sets as a significant improvement over previous RNA-seq based treatments of the basidiomycete life cycle and we anticipate that beyond starvation, photobiology and mycelial differentiation, these data will contribute to a better understanding of the fruiting process. We also expect the data to provide further insights into the photobiology of basidiomycetes and open the way to comparative analyses of light responses of basidiomycetes with those of the Ascomycota (e.g. *N. crassa*), in which the process is much better understood (114).

These expression data, along with the new and a previously published *C. cinerea* genome, annotation and transcriptomes, were integrated into a new website (mushroomdb.brc.hu), which we plan to update regularly to provide a reference resource for the mushroom science community for future functional analyses. The website is centered around *C. cinerea*, as the currently best-known mushroom-forming model species, but includes data from several industrially relevant and commercially produced mushroom-forming species too.

Some limitations resulting from the employed technologies should be mentioned. While we expect the vast majority of genes to be correctly annotated, the annotation of some genes (e.g., lowly expressed long genes) or features (e.g., TSS) should be improved further. Given that we used stringent filtering criteria to assemble the gene catalog, it is possible that future studies will discover that some, albeit likely not many, further genes can be found in *C. cinerea*. While we validated the TTS positions of the gene models by Quantseq, the TSS were not experimentally checked. This may lead to some imprecisions in the TSS positions, which may be remedied by assays such as SAGE, CAGE (58, 115). The expression data generated in this study allowed the identification of genes responsive for light, starvation and tissue type. However, given the diversity of conditions, we envision, these data could be used for addressing more complex questions on gene expression, such as co-expression patterns or global regulatory networks. The integration of the data into a dedicated website (mushroomdb.brc.hu) is a step in this direction, whereas global gene regulatory network reconstruction is a future challenge. An exciting next step towards network-based approaches is exploring patterns of cis-regulation in *C. cinerea*, by high-throughput functional assays (e.g., ATAC-, DAP-, or ChIP-seq), which will allow deeper understanding of the mechanism of gene expression regulation under diverse conditions. Regulatory interactions could also be interrogated by directed perturbation of the network by reverse genetics, as shown recently by analyses of the *cre1, snb1* and *nsdD1 and nsdD2* genes (79, 94, 116). Another possible extension of this work is improving the assemblies, annotations and gene functions in other non-model basidiomycetes too, to enable rigorous comparative analysis of gene model and expression patterns across species.

Taken together, this study has improved the genome assembly, annotation as well as expanded our functional knowledge of protein-coding genes of *C. cinerea*. With these improvements, *C. cinerea* emerges as potentially the most comprehensively annotated species among the Agaricomycetes. We anticipate this work will serve as a foundational resource for further studies, particularly those in which genome integrity and accurate gene model prediction are essential, and will facilitate further research on mushroom-forming and lignocellulose-degrading Agaricomycetes as well as facilitate functional genomic comparisons across the fungal kingdom.

## Materials and methods

### Growth conditions

For all experiments we used the *C. cinerea* #326 Amut1Bmut1 PABA-strain (FGSC#25122), which carries mutations in both mating type factors that overcomes the self-incompatibility of monokaryotic strains (117). High molecular weight DNA for PacBio sequencing was extracted from mycelium samples grown in liquid YMG (0.4 % yeast extract, 1% malt extract, 0.4% glucose medium at 28 °C without shaking using the Blood and Cell Culture DNA Maxi Kit (Qiagen, catalog number: 13362) following the manufacturer’s instructions. RNA for Nanopore and IsoSeq was extracted from mycelium and fruiting body tissues using Quick-RNA Miniprep kit (Zymo Research, USA, catalog number: R1054), following the manufacturer’s instructions. The same kit was used for RNA extraction for gene expression profiling. Supplementary Table 5 provides detailed growth conditions and protocols for gene expression profiling experiments.

### Genome sequencing and assembly

Approximately 10 µg of genomic DNA was sheared to 30-50 kb using the Megaruptor 3 (Diagenode). The sheared DNA was treated with exonuclease to remove single-stranded ends, DNA damage repair enzyme mix, end-repair A-tailing mix, and ligated with overhang adapters using SMRTbell Express Template Prep Kit 2.0 (PacBio) and purified with AMPure PB Beads (PacBio). Individual libraries were size-selected (20 kb) using the 0.75% agarose gel cassettes with Marker S1 and High Pass protocol on the Blue Pippin instrument (Sage Science). PacBio sequencing primer was then annealed to the SMRTbell template library and sequencing polymerase was bound with a Sequel II Binding kit 2.0. The prepared SMRTbell template libraries were then sequenced on a Pacific Biosystems Sequel II sequencer using tbd-sample dependent sequencing primer, 8M v1 SMRT cells, and Version 2.0 sequencing chemistry with 1x1800 sequencing movie run times.

Filtered subread data was processed with the JGI QC pipeline to remove artifacts.

The mitochondrial genome was assembled separately with the CCS reads using an in-house tool, used to filter the CCS reads, and polished with two rounds of Racon version 1.4.13 (118) racon [-u -t 36] (118). The mitochondria-filtered CCS reads were then assembled with Flye version 2.8.1-b1676 (119) [-g 40M --asm-coverage 50 -t 32 --pacbio-hifi] to generate an assembly and polished with two rounds of RACON version 1.4.13 (118) racon [-u -t 36].

Ribosomal DNA was assembled separately from a subset of CCS reads identified using kmer matching against the UNITE database with BBTools version 38.79 (120) bbduk.sh [k=31 mm=f mkf=0.05 ow=true]. Matching reads were subsampled to 600000bp with BBTools (120) reformat.sh [sampleseed=1 samplebasestarget=600000 prioritizelength=t ow=true] and assembled with Flye version 2.8.1 --py37h4bd9754_0 (119) [--hifi-error 0.01 --meta --keep-haplotypes --genome-size 12K -t 16 --pacbio-hifi] and polished with two rounds of RACON version 1.4.13 (118) racon [-u -t 36].

Eukaryotic internal transcribed spacer (ITS) was identified and extracted from the rDNA assembly using ITSx version quay.iobiocontainersitsx1.1b--2 (121) [--detailed_results T -- anchor 100 --cpu 16 -t F -o ITS]. Results were used to orient and trim the rDNA contig to 100bp SSU - 1Kb LSU.

### PacBio Iso-seq library preparation, sequencing and read processing

Full-length cDNA was synthesized using template switching technology with NEBNext Single Cell/Low Input cDNA Synthesis & Amplification Module kit. The first-strand cDNA was amplified and multiplexed with NEBNext High-Fidelity 2X PCR Master Mix using Barcoded cDNA PCR primers. The amplified cDNA was purified using 1.3X ProNex beads. Similar sizes were pooled at equimolar ratios in a designated Degree-of-Pool using PacBio Multiplexing Calculator. The pooled samples were end-repaired, A-tailed and ligated with overhang non-barcoded adaptors using SMRTbell Express 2.0 kit. PacBio Sequencing primer was then annealed to the SMRTbell template library and sequencing polymerase was bound to them using Sequel II Binding kit 2.0. The prepared SMRTbell template libraries were then sequenced on a Pacific Biosystems’ Sequel II sequencer using sequencing primer, 8M v1 SMRT cells, and Version 2.0 sequencing chemistry with 1x1800 sequencing movie run times.

PacBio subreads were used as an input to generate the polished Circular Consensus Sequence by pbccs (version 4). In pbccs the accuracy rate and the minimum pass number were set to 98% and 3 ( ccs --min-passes 3 --min-snr 4 --max-length 21000 --min-length 10 --min-rq 0.98). For the classification of the css reads into full length category according to the presence of the 5’ primer, 3’ primer and poly(A) tail lima was used. For the further classification only, the full-length reads were kept. In the further steps isoseq3 refine function was used to remove poly(A) tails and cluster function to cluster all the full length css reads.

### Oxford nanopore cDNA library preparation, sequencing and read processing

From each of the five purified total RNA samples, 2 µg of RNA was used to prepare full-length cDNA libraries using the TeloPrime Full-Length cDNA Amplification Kit V2 (Lexogen, Vienna) protocol. Endpoint PCR was performed using the same kit with 12 to 18 cycles. The concentration and integrity of the samples were determined using a Qubit 2.0 fluorometer with the Qubit (ds)DNA HS Assay Kit (Thermo Fisher Scientific) and the Bioanalyzer 2100 (Agilent Technologies) with the DNA 7500 chip. The amplified cDNA samples were barcoded using the Oxford Nanopore Native Barcoding Expansion Kit (EXP-NBD104). The proportionally mixed barcoded samples were used to prepare libraries with the Ligation Sequencing Kit (SQK-LSK109, Oxford Nanopore Technologies), which were then sequenced on two MinION flow cells (FLO-MIN106 R9.4.1, Oxford Nanopore Technologies). Base calling and demultiplexing were performed using Guppy v3.2.4. Only reads with a Q value ≥7 were used in further steps. The base called reads were adapter trimmed and oriented using Pychopper v2.3.1 (122). Any reads shorter than 50 bp were excluded. Errors in ONT reads were polished using LoRDEC v0.9 (123). For the polishing process BBTools reformat.sh (120) was used to randomly sample approximately 20M R1 Illumina reads from a previous dataset (24). To remove redundant or poor-quality reads from the sampled dataset prior to the polishing step, reads were further sampled using ORNA (124). To evaluate the indel and mismatch ratios of the long reads assembled with CopciAB V2, the functions stats_from_bam and summary_from_stats from P (125) were used, which output summary statistics for each read.

### Quantseq library preparation, sequencing

The libraries were generated using the QuantSeq 3’ mRNA-Seq Library Prep Kit FWD for Illumina (Lexogen, Vienna). To begin, 100 ng of total RNA was used as input for first strand cDNA generation with an oligo-dT primer, followed by RNA removal. Next, second strand synthesis was initiated with random priming and the resulting products were purified with magnetic beads. Finally, the libraries were amplified and barcoded using PCR. The Agilent 4200 TapeStation (Agilent) was used to assess all libraries for the formation of adapter dimers during PCR. The QuantSeq libraries were sequenced on the Illumina NextSeq550 platform, producing 75 bp single-end reads.

### Genome comparison, structural variation identification

To study the differences in chromosomes (C13) three genomes were compared using the dnadiff function of the MUMmer v4.0 package (126) with CopciAB V2 as a reference. We identified structural variations (SVs) between the genomes using SyRI (127). For this purpose, chromosome-level alignment was performed between *C. cinerea* Okayama (reference) and CopciAB V2 (query) genomes, as well as between CopciAB V2 (reference) and ONT-based (query) genomes using Nucmer from the MUMmer package. The alignments were used as input for SyRI to identify the genomic rearrangements and local sequence differences. The output of SyRI was plotted using plotsr (128).

### Determination of indel, SNP variation between the CopciAB V2 and Copci ONT genomes

Genomic Illumina reads (SRR10162428) (29) were trimmed with fastp v0.21.0 (129) and then mapped to the CopciAB V2 and CopciAB ONT genomes with bwa v0.7.17-r1198-dirty (130). After sorting and preprocessing the bam alignment files with samtools v1.13 (131) and picard package AddOrReplaceReadGroups v2.18.14-SNAPSHOT function (132), the indels and SNPs were called using GATK3 (133).

### Long-read based gene model prediction

During gene model prediction, the main objective was to integrate data from all available genomes and associated annotations into a more complete long-read based annotation. To accomplish this we iteratively utilized the Illumina (28), ONT (29) and the new PacBio genome. During each iteration, we incorporated manually curated gene models from the reference genome and newly created ONT based gene models into the resulting annotation. In the final polishing step, we integrated the information carried by the PacBio Isoseq reads into the reference annotation, resulting in the final form.

In detail, we performed the following steps. Firstly, we separately mapped the polished ONT reads from five different samples to the CopciAB V1 genome (28) using DeSALT (134). We chose this aligner due to its higher observed precision in *de novo* mapping of transcripts with shorter exons compared to other long read aligners (minimap2 v2.24-r1122 (135), magicblast: v1.5.0 (136), and GraphMap v0.6.3 (137)). The aligned reads were processed using samclip (https://github.com/tseemann/samclip) to remove long soft clip sequences. The resulting trimmed alignments were then utilized to predict gene models with the TAMA software package (138). Using TAMA, we initially collapsed mapped reads by samples [tama_collapse.py -s $inputBAMfile -b BAM -f $refgenomePath -p $outTag -x capped -icm ident_map -a 50 -z 50]. Subsequently, we filtered out collapsed reads with low read support (at least 2 reads in at least 1 sample) [tama_remove_single_read_models_levels.py -b$annotationBedFile -r $readSupportFile -l transcript -k remove_multi -o $outputPrefix]. Finally, we merged the collapsed reads from the different samples [tama_merge.py -f $fileList -p $outputPrefix -a 50 -z 50].

It was observed that TAMA often fails to separate overlapping neighboring gene models. To address this issue, we reviewed the length of each TAMA-suggested transcript isoform for each gene model and. To eliminate exceptionally long, possible erroneous isoforms, we filtered out any isoforms with a length twice as larger as the geometric mean of the gene’s isoforms. The remaining isoforms were clustered based on their overlap ratio. If the pairwise overlap was less than 50%, the isoforms were reassigned to a separate gene model represented by the isoform with the highest read support. To preserve all the data, the previously filtered out, especially long isoforms were later added back to the gene cluster with the highest read support. In a further step, ORFik (139) was then utilized to predict coding DNA sites (CDSs) with a minimum length of 30 amino acids on the gene/transcript models. Transcript models without CDS predictions were omitted from further analysis.

To identify the primary transcript isoform of the gene for further manual curation, certain filter criteria were required to be met. Firstly, a transcript must be longer than the 80% of the geometric mean of all transcripts in the cluster. Secondly, if a transcript has at least 4 times higher expression than the second highest expressed transcript, it becomes the primary isoform. Otherwise, the longest isoform is selected. The primary isoforms selected were manually checked. To aid in the curation process, Illumina reads (24) were aligned to the reference genome using the STAR v2.6.1a_08-27 aligner. Non-canonical splice sites, splice junctions (SJ) with low Illumina read support (≤5 reads), or alignment errors (indel, mismatch) around the SJ were given special attention and were manually checked and corrected if necessary. At this point, any gene clusters that were removed due to the above 80% filter were restored. Also, partial primary isoforms were corrected to accurately reflect the CDS. Gene models not present in the primary isoforms, but present in the CopciAB V1 annotations were incorporated.

The primary isoforms defined in the CopciAB V1 genome were mapped onto the ONT-based genome (29) using DeSALT. We included gene models that were exclusively present in the ONT-based genome to the primary isoform set. In case when there was a match between the gene models of the ONT-based genome and CopciAB V1 genome, we gave priority to the ones that originated from the ONT-based genome over the CopciAB V1. We utilized SQANTI3 (140) to identify overlapping gene models. We then used DeSALT to map the extended primary isoform set onto the PacBio-based genome. To detect alignment errors for manual curation, we aligned Illumina reads as previously mentioned.

To further improve annotation, we incorporated PacBio Isoseq reads. We aligned ONT and PacBio reads to the reference genome using minimap2 with the assistance of reference annotation. We excluded long reads that had a perfect SJ match with the reference annotation and only considered them as read support for the corresponding gene. The remaining long reads were utilized for an additional round of transcript model prediction with TAMA.

The second round of TAMA transcript/gene models were compared with the corrected extended primary isoform. If the length of the CDS encoded by the new gene/transcript model was longer than the counterpart in the reference genome, then the new gene/transcript model was accepted. If the CDS was identical to a CopciAB V1 (28) or the ONT-based (29) gene in the reference genome then, it was also replaced with the new gene model. If it was missing, it was transferred to the corrected extended primary isoform. The gene models that were taken from the new TAMA run underwent manual checking before being accepted. To support the representation of the long gene models with low expression in the long-read based annotation, the long genes with one PacBio Isoseq read support were also accepted after consideration.

Transcript isoforms were accepted if they had a read support of at least 10 reads and accounted for at least 10% of the gene’s total read support, which includes both ONT and PacBio Isoseq reads.

To identify falsely predicted gene models, we first identified genes with extremely low expression. We collected 165 RNA-seq samples from five studies, each with at least two biological replicates, including two published (24, 112), one published with the current work (Quantseq data), and two currently unpublished studies (CPM values available on the mushroomdb.brc.hu webpage), For each gene in each sample, we calculated the mean count per million (CPM). Next, we focused on the highest possible mean CPM of the gene reached per study. We marked genes whose maximum expression fell in the lowest 10% of the expression maxima in all five studies as low expressed genes. Low expressed genes without similarity to proteins outside of *C. cinerea* (mmseqs, evalue ≤1e-10, qcov ≥0.5), lacking Interpro annotation and long read support, were filtered out from the reference annotation. The completeness of the new annotation was assessed using BUSCO v3.0.2 with basidiomycota_db10 database (141).

### Determination of isoforms

SUPPA2 (142) was used for splice isoform classification. To improve the detection of intron retention events (RI), the program was executed in two rounds. Initially, the default parameters were utilized, followed by a more permissive second run using the parameters -b V -t 50. The RI events from the first run were supplemented by the RI events from the second run.

### Identification of the conserved genomic elements

To identify the conserved genomic elements around transcription start site (TSS), we retrieved sequences of ± 50 nt flanking the TSS position of the long-read-based gene models with 5’UTR region. For the sequence context of the transcription initiation site (TIS), we retrieved sequences of ± 12 nt flanking the ATG position of the long-read-based gene models with a minimum 12 nt long 5’UTR region. In the case of the splice site, 6 nt from the 5’ end and 3 nt from the 3’ end of the intron sequences were retrived and used to identify 5’ss and 3’ss. Finally, in the case of the transcriptional termination site (TTS), ± 50 nt of sequence flanking the TTS were recovered from long-read-based gene models with 3’UTR region. The sequences obtained were analyzed using ggseqlogo (143). For the identification of the branch point (BP) the branchpointer v1.14 R package was used (144).

### PASs, PACs identification. Relative PAC usage determination

For polyadenylation site (PAS) identification, the DPAC tool (145) was used all corresponding QuantSeq reads. Briefly, the QuantSeq reads were adapter and poly(A) trimmed and the length of the poly(A) tail was recorded. Reads with at least 15 A’s were mapped to the reference genome. The 3’ end of all reads was used to identify the position of PASs in the reference genome. PASs with at least 20 reads, each with a poly(A) tail of at least 15 A’s, were retained. PASs were further filtered for internal priming and template switching by counting the number of A’s in the reference genome immediately downstream of the identified PAS. If at least a 4 nt long poly(A) stretch was found immediately downstream of the PAS or, ≥70% of the As were in a 10 nt long downstream window, the PAS was filtered out from the further analysis.

The PAS were further clustered into polyadenylation site clusters (PAC) by allowing a maximum intra-cluster distance of 10 nt. Within a PAC, the PAS with the highest read support serves as the representative PAS of the cluster and lends its read support and genomic coordinate to the PAC (Figure 3/A). This common coordinate was used to assign the PACs to the different entries of the genome annotation and to identify the associated gene models by assigning them to the TTS of the closest gene. To further reduce the negative effects of internal priming and template switching on PAC identification, we only accepted those PACs which are located in the 3’UTR of the associated gene and in the downstream intergenic space, or in the genomic region of the antisense gene within 1,000 nt of the TTS of the associated gene (Figure 3/B). A distinction was made between PACs based on read support. The PACs of the gene with the highest read support, the highest representative PAS read support, were considered the best supported PAC.

PAC usage across biological conditions was defined by sample-wise counting of mapped 3’ ends of the previously trimmed QuantSeq reads on the defined PACs. PAC read counts were normalized to CPM. The normalized counts were pooled by conditions for increased sensitivity.

To determine the evenness value of APA genes with ≥10 CPM, the vegan R package (146) was used to calculate the Shannon diversity index for the PAC sites and dividing by the logarithmic value of the number of PAC sites. The line graph showing the correlation between the value of the evenness of the APA gene PAC sites and the value of gene expression was generated using ggplot2 (147). The lines were smoothed using a generalized additive model (GAM) (148).

### Gene expression analysis

The new *C. cinerea* Amut1Bmut1 #326 genome and gene model prediction were used as a reference for gene expression analysis. The quality of the raw and trimmed QuantSeq reads was assessed using the FastQC (149) and MultiQC (150) tools. The raw reads were trimmed using the BBduk.sh script from BBtools 38.92 (120) and then mapped to the reference genome using STAR version 2.6.1a_08-27 (151). The trimming and mapping parameters were set according to the manufacturer’s recommendations (https://www.lexogen.com/quantseq-data-analysis/). The number of reads overlapping the transcript annotations was determined using the featureCounts function of the Rsubread package version 2.6.4 (152). For differential gene expression analysis, the edgeR version 3.34.0 (153) and limma version 3.48.3 (154) packages were used. To minimize the number of misidentified differentially expressed genes (DEGs), only genes whose expression reached at least 1 CPM in at least one experiment were included in the analysis. Genes with a fold change ≥|2| and a Benjamini-Hochberg (BH) adjusted *p* value ≤0.05 were considered as significantly differentially expressed DEGs. To further minimize the number of misidentified DEGs, we only included those DEGs in the further analysis that had an average of at least 2 CPM in at least one of the conditions during comparison. The ComplexHeatmap R library (155) was used to generate expression heatmaps.

### Functional annotation and GO enrichment analysis

InterPro (IPR) and GO functional annotations were defined using InterProScan version 93 (156). Putative CAZymes were identified using dbCAN (157) and further categorized based on their substrate specificity as described in previous publications (158). Based on the substrate specificity information, the groups of plant cell wall degrading CAZymes (PCWDEs) and fungal cell wall degrading CAZymes (FCWDEs) were defined (available at mushroomdb.brc.hu). The gene sets for transcription factors (TFs) and transporters were defined using InterPro entry information and evidence from previous publications (available at mushroomdb.brc.hu). GO enrichment analysis was performed using topGO version 2.40.0 (159) in conjunction with GO.db version 3.17.0 (160). The significantly enriched GO terms were plotted with the R package ggplot2 (147).

## Acknowledgements

The authors appreciate valuable discussions and suggestions on Nanopore sequencing from Zsolt Boldogkoi and Dora Tombacz (University of Szeged).

## Data availability

## References

1. Floudas, D., Binder, M., Riley, R., Barry, K., Blanchette, R., Henrissat, B., Martínez, A.T., Otillar, R., Spatafora, J.W., Yadav, J.S., et al. (2012) The Paleozoic Origin of Enzymatic Lignin Decomposition Reconstructed from 31 Fungal Genomes. Science, 336, 1715– 1719.

2. Virágh, M., Merényi, Z., Csernetics, Á., Földi, C., Sahu, N., Liu, X.-B., Hibbett, D.S. and Nagy, L.G. (2021) Evolutionary Morphogenesis of Sexual Fruiting Bodies in Basidiomycota: Toward a New Evo-Devo Synthesis. Microbiol. Mol. Biol. Rev., 86, e00019–21.

3. Grigoriev, I.V., Nikitin, R., Haridas, S., Kuo, A., Ohm, R., Otillar, R., Riley, R., Salamov, A., Zhao, X., Korzeniewski, F., et al. (2014) MycoCosm portal: Gearing up for 1000 fungal genomes. Nucleic Acids Res., 42, 699–704.

4. Kües, U. and Navarro-González, M. (2015) How do Agaricomycetes shape their fruiting bodies? 1. Morphological aspects of development. Fungal Biol. Rev., 29, 63–97.

5. Yang, C., Ma, L., Xiao, D., Liu, X., Jiang, X., Ying, Z. and Lin, Y. (2021) Chromosome-scale assembly of the Sparassis latifolia genome obtained using long-read and Hi-C sequencing. G3 Genes Genomes Genet., 11.

6. Yu, H., Zhang, L., Shang, X., Peng, B., Li, Y., Xiao, S., Tan, Q. and Fu, Y. (2022) Chromosomal genome and population genetic analyses to reveal genetic architecture, breeding history and genes related to cadmium accumulation in Lentinula edodes. BMC Genomics, 23, 1–14.

7. Jo, I.H., Kim, J., An, H., Lee, H.Y., So, Y.S., Ryu, H., Sung, G.H., Shim, D. and Chung, J.W. (2022) Pseudo-Chromosomal Genome Assembly in Combination with Comprehensive Transcriptome Analysis in Agaricus bisporus Strain KMCC00540 Reveals Mechanical Stimulus Responsive Genes Associated with Browning Effect. J. Fungi, 8, 1–15.

8. Zhu, Y., Xu, J., Sun, C., Zhou, S., Xu, H., Nelson, D.R., Qian, J., Song, J., Luo, H., Xiang, L., et al. (2015) Chromosome-level genome map provides insights into diverse defense mechanisms in the medicinal fungus Ganoderma sinense. Sci. Rep., 5, 1–14.

9. Yu, H., Zhang, M., Sun, Y., Li, Q., Liu, J., Song, C., Shang, X., Tan, Q., Zhang, L. and Yu, H. (2022) Whole-genome sequence of a high-temperature edible mushroom Pleurotus giganteus (zhudugu). Front. Microbiol., 13, 1–11.

10. Sonnenberg, A.S.M., Sedaghat-Telgerd, N., Lavrijssen, B., Ohm, R.A., Hendrickx, P.M., Scholtmeijer, K., Baars, J.J.P. and van Peer, A. (2020) Telomere-to-telomere assembled and centromere annotated genomes of the two main subspecies of the button mushroom Agaricus bisporus reveal especially polymorphic chromosome ends. Sci. Rep., 10, 1– 15.

11. Lee, Y.Y., de Ulzurrun, G.V.D., Schwarz, E.M., Stajich, J.E. and Hsueh, Y.P. (2021) Genome sequence of the oyster mushroom Pleurotus ostreatus strain PC9. G3 Genes Genomes Genet., 11.

12. Galagan, J.E., Henn, M.R., Ma, L.J., Cuomo, C.A. and Birren, B. (2005) Genomics of the fungal kingdom: Insights into eukaryotic biology. Genome Res., 15, 1620–1631.

13. Wang, L., Jiang, N., Wang, L., Fang, O., Leach, L.J., Hu, X. and Luo, Z. (2014) 3′ Untranslated Regions Mediate Transcriptional Interference between Convergent Genes Both Locally and Ectopically in Saccharomyces cerevisiae. PLoS Genet., 10, 1–12.

14. Gilet, J., Conte, R., Torchet, C., Benard, L. and Lafontaine, I. (2019) Additional Layer of Regulation via Convergent Gene Orientation in Yeasts. Mol. Biol. Evol., 37, 365–378.

15. Gordon, S.P., Tseng, E., Salamov, A., Zhang, J., Meng, X., Zhao, Z., Kang, D., Underwood, J., Grigoriev, I.V., Figueroa, M., et al. (2015) Widespread polycistronic transcripts in fungi revealed by single-molecule mRNA sequencing. PLoS ONE, 10, 1–15.

16. Lu, P., Chen, D., Qi, Z., Wang, H., Chen, Y., Wang, Q., Jiang, C., Xu, J.R. and Liu, H. (2022) Landscape and regulation of alternative splicing and alternative polyadenylation in a plant pathogenic fungus. New Phytol., 235, 674–689.

17. Leppek, K., Das, R. and Barna, M. (2018) Functional 5′ UTR mRNA structures in eukaryotic translation regulation and how to find them. Nat. Rev. Mol. Cell Biol., 19, 158–174.

18. Araujo, P.R., Yoon, K., Ko, D., Smith, A.D., Qiao, M., Suresh, U., Burns, S.C. and Penalva, L.O.F. (2012) Before it gets started: Regulating translation at the 5′ UTR. Comp. Funct. Genomics, 2012.

19. Mayr, C. (2017) Regulation by 3′-Untranslated Regions. 10.1146/annurev-genet-120116-024704.

20. Ransom, B., Goldman, S.A., Meldolesi, J., Zhou, L., Murai, K.K., Harris, K.M., Mccarthy, K.D., Li, N., Doyle, R.T., Haydon, P.G., et al. (2008) Proliferating Cells Express mRNAs with Shortened 3′ Untranslated Regions and Fewer MicroRNA Target Sites. Science, 320, 1643–1647.

21. Mayr, C. and Bartel, D.P. (2009) Widespread Shortening of 3′UTRs by Alternative Cleavage and Polyadenylation Activates Oncogenes in Cancer Cells. Cell, 138, 673–684.

22. Tushev, G., Glock, C., Heumüller, M., Biever, A., Jovanovic, M. and Schuman, E.M. (2018) Alternative 3′ UTRs Modify the Localization, Regulatory Potential, Stability, and Plasticity of mRNAs in Neuronal Compartments. Neuron, 98, 495–511.e6.

23. Kües, U. (2000) Life History and Developmental Processes in the Basidiomycete Coprinus cinereus. Microbiol. Mol. Biol. Rev., 64, 316–353.

24. Krizsan, K., Almasi, E., Merenyi, Z., Sahu, N., Viragh, M., Koszo, T., Mondo, S., Kiss, B., Balint, B., Kues, U., et al. (2019) Transcriptomic atlas of mushroom development highlights an independent origin of complex multicellularity in fungi. Proc. Natl. Acad. Sci. U. S. A., 10.1073/pnas.1817822116.

25. Virágh, M., Merényi, Z., Csernetics, Á., Földi, C., Sahu, N., Liu, X. and Hibbett, D.S. Evolutionary morphogenesis of sexual fruiting bodies in Basidiomycota: toward a new evo-devo synthesis Máté Virágh. 26.

26. Stajich, J.E., Wilke, S.K., Ahrén, D., Au, C.H., Birren, B.W., Borodovsky, M., Burns, C., Canbäck, B., Casselton, L.A., Cheng, C.K., et al. (2010) Insights into evolution of multicellular fungi from the assembled chromosomes of the mushroom Coprinopsis cinerea (Coprinus cinereus). Proc. Natl. Acad. Sci. U. S. A., 107, 11889–11894.

27. Ruiz-Dueñas, F.J., Barrasa, J.M., Sánchez-García, M., Camarero, S., Miyauchi, S., Serrano, A., Linde, D., Babiker, R., Drula, E., Ayuso-Fernández, I., et al. (2021) Genomic Analysis Enlightens Agaricales Lifestyle Evolution and Increasing Peroxidase Diversity. Mol. Biol. Evol., 38, 1428–1446.

28. Muraguchi, H., Umezawa, K., Niikura, M., Yoshida, M., Kozaki, T., Ishii, K., Sakai, K., Shimizu, M., Nakahori, K., Sakamoto, Y., et al. (2015) Strand-specific RNA-seq analyses of fruiting body development in Coprinopsis cinerea. PLoS ONE, 10, 1–23.

29. Xie, Y., Zhong, Y., Chang, J. and Kwan, H.S. (2021) Chromosome-level de novo assembly of Coprinopsis cinerea A43mut B43mut pab1-1 #326 and genetic variant identification of mutants using Nanopore MinION sequencing. Fungal Genet. Biol., 146, 103485.

30. Pukkila, P.J. (2011) Coprinopsis cinerea. Curr. Biol., 21, R616–R617.

31. Sakamoto, Y. (2018) Influences of environmental factors on fruiting body induction, development and maturation in mushroom-forming fungi. Fungal Biol. Rev., 32, 236– 248.

32. Ustianenko, D., Weyn-Vanhentenryck, S.M. and Zhang, C. (2017) Microexons: discovery, regulation, and function. Wiley Interdiscip. Rev. RNA, 8.

33. Li, Y.I., Sanchez-Pulido, L., Haerty, W. and Ponting, C.P. (2015) RBFOX and PTBP1 proteins regulate the alternative splicing of micro-exons in human brain transcripts. Genome Res., 25, 1–13.

34. Irimia, M., Weatheritt, R.J., Ellis, J.D., Parikshak, N.N., Gonatopoulos-Pournatzis, T., Babor, M., Quesnel-Vallières, M., Tapial, J., Raj, B., O’Hanlon, D., et al. (2014) A highly conserved program of neuronal microexons is misregulated in autistic brains. Cell, 159, 1511–1523.

35. Cooper, T.A. and Ordahl, C.P. (1985) A single cardiac troponin T gene generates embryonic and adult isoforms via developmentally regulated alternate splicing. J. Biol. Chem., 260, 11140–11148.

36. Parada, G.E., Munita, R., Georgakopoulos-Soares, I., Fernandes, H.J.R., Kedlian, V.R., Metzakopian, E., Andres, M.E., Miska, E.A. and Hemberg, M. (2021) MicroExonator enables systematic discovery and quantification of microexons across mouse embryonic development. Genome Biol., 22, 43.

37. McAllister, L., Rehm, E.J., Goodman, C.S. and Zinn, K. (1992) Alternative splicing of micro-exons creates multiple forms of the insect cell adhesion molecule fasciclin I. J. Neurosci., 12, 895–905.

38. Chang, L.W., Tseng, I.C., Wang, L.H. and Sun, Y.H. (2020) Isoform-specific functions of an evolutionarily conserved 3 bp micro-exon alternatively spliced from another exon in Drosophila homothorax gene. Sci. Rep., 10, 1–13.

39. Guo, L. and Liu, C.M. (2015) A single-nucleotide exon found in Arabidopsis. Sci. Rep., 5, 1–5.

40. Song, Q., Lv, F., Qamar, M.T.U., Xing, F., Zhou, R., Li, H. and Chen, L.L. (2019) Identification and analysis of micro-exon genes in the rice genome. Int. J. Mol. Sci., 20, 1–14.

41. Wang, K., Wang, D., Zheng, X., Qin, A., Zhou, J., Guo, B., Chen, Y., Wen, X., Ye, W., Zhou, Y., et al. (2019) Multi-strategic RNA-seq analysis reveals a high-resolution transcriptional landscape in cotton. Nat. Commun., 10.

42. Schotanus, K., Soyer, J.L., Connolly, L.R., Grandaubert, J., Happel, P., Smith, K.M., Freitag, M. and Stukenbrock, E.H. (2015) Histone modifications rather than the novel regional centromeres of Zymoseptoria tritici distinguish core and accessory chromosomes. Epigenetics Chromatin, 8, 41.

43. Sepsiova, R., Necasova, I., Willcox, S., Prochazkova, K., Gorilak, P., Nosek, J., Hofr, C., Griffith, J.D. and Tomaska, L. (2016) Evolution of Telomeres in Schizosaccharomyces pombe and Its Possible Relationship to the Diversification of Telomere Binding Proteins. PLoS ONE, 11, e0154225.

44. Wellinger, R.J. and Zakian, V.A. (2012) Everything You Ever Wanted to Know About Saccharomyces cerevisiae Telomeres: Beginning to End. Genetics, 191, 1073–1105.

45. de Lange, T. (2004) T-loops and the origin of telomeres. Nat. Rev. Mol. Cell Biol., 5, 323– 329.

46. Swapna, G., Yu, E.Y. and Lue, N.F. (2018) Single telomere length analysis in Ustilago maydis, a high-resolution tool for examining fungal telomere length distribution and C- strand 5’-end processing. Microb. Cell, 5, 393–403.

47. Heinzelmann, R., Rigling, D., Sipos, G., Münsterkötter, M. and Croll, D. (2020) Chromosomal assembly and analyses of genome-wide recombination rates in the forest pathogenic fungus Armillaria ostoyae. Heredity, 124, 699–713.

48. Pérez, G., Pangilinan, J., Pisabarro, A.G. and Ramírez, L. (2009) Telomere organization in the ligninolytic basidiomycete pleurotus ostreatus. Appl. Environ. Microbiol., 75, 1427– 1436.

49. Saud, Z., Kortsinoglou, A.M., Kouvelis, V.N. and Butt, T.M. (2021) Telomere length de novo assembly of all 7 chromosomes and mitogenome sequencing of the model entomopathogenic fungus, Metarhizium brunneum, by means of a novel assembly pipeline. BMC Genomics, 22, 1–15.

50. Ke, H.-M., Lee, H.-H., Lin, C.-Y.I., Liu, Y.-C., Lu, M.R., Hsieh, J.-W.A., Chang, C.-C., Wu, P.-H., Lu, M.J., Li, J.-Y., et al. (2020) Mycena genomes resolve the evolution of fungal bioluminescence. Proc. Natl. Acad. Sci., 117, 31267–31277.

51. Jenjaroenpun, P., Wongsurawat, T., Pereira, R., Patumcharoenpol, P., Ussery, D.W., Nielsen, J. and Nookaew, I. (2018) Complete genomic and transcriptional landscape analysis using third-generation sequencing: a case study of Saccharomyces cerevisiae CEN.PK113-7D. Nucleic Acids Res., 10.1093/nar/gky014.

52. Weirather, J.L., de Cesare, M., Wang, Y., Piazza, P., Sebastiano, V., Wang, X.-J., Buck, D. and Au, K.F. (2017) Comprehensive comparison of Pacific Biosciences and Oxford Nanopore Technologies and their applications to transcriptome analysis. F1000Research, 6, 100.

53. Sessegolo, C., Cruaud, C., Da Silva, C., Cologne, A., Dubarry, M., Derrien, T., Lacroix, V. and Aury, J.M. (2019) Transcriptome profiling of mouse samples using nanopore sequencing of cDNA and RNA molecules. Sci. Rep., 9, 1–12.

54. Balázs, Z., Tombácz, D., Csabai, Z., Moldován, N., Snyder, M. and Boldogkoi, Z. (2019) Template-switching artifacts resemble alternative polyadenylation. BMC Genomics, 20, 1–10.

55. Doddapaneni, H., Chakraborty, R. and Yadav, J.S. (2005) Genome-wide structural and evolutionary analysis of the P450 monooxygenase genes (P450ome) in the white rot fungus Phanerochaete chrysosporium : Evidence for gene duplications and extensive gene clustering. BMC Genomics, 6, 92.

56. Wallace, E.W.J., Maufrais, C., Sales-Lee, J., Tuck, L.R., de Oliveira, L., Feuerbach, F., Moyrand, F., Natarajan, P., Madhani, H.D. and Janbon, G. (2020) Quantitative global studies reveal differential translational control by start codon context across the fungal kingdom. Nucleic Acids Res., 48, 2312–2331.

57. Zhang, Z. and Dietrich, F.S. (2005) Mapping of transcription start sites in Saccharomyces cerevisiae using 5′ SAGE. Nucleic Acids Res., 33, 2838–2851.

58. Hashimoto, S.I., Suzuki, Y., Kasai, Y., Morohoshi, K., Yamada, T., Sese, J., Morishita, S., Sugano, S. and Matsushima, K. (2004) 5′-end SAGE for the analysis of transcriptional start sites. Nat. Biotechnol., 22, 1146–1149.

59. Sibthorp, C., Wu, H., Cowley, G., Wong, P.W.H., Palaima, P., Morozov, I.Y., Weedall, G.D. and Caddick, M.X. (2013) Transcriptome analysis of the filamentous fungus Aspergillus nidulans directed to the global identification of promoters. BMC Genomics, 14.

60. Li, H., Hou, J., Bai, L., Hu, C., Tong, P., Kang, Y., Zhao, X. and Shao, Z. (2015) Genome-wide analysis of core promoter structures in Schizosaccharomyces pombe with DeepCAGE. RNA Biol, 12, 525–537.

61. Kupfer, D.M., Drabenstot, S.D., Buchanan, K.L., Lai, H., Zhu, H., Dyer, D.W., Roe, B.A. and Murphy, J.W. (2004) Introns and splicing elements of five diverse fungi. Eukaryot. Cell, 3, 1088–1100.

62. Mata, J. (2013) Genome-wide mapping of polyadenylation sites in fission yeast reveals widespread alternative polyadenylation. RNA Biol., 10, 1407–1414.

63. Liu, X., Hoque, M., Larochelle, M., Lemay, J.F., Yurko, N., Manley, J.L., Bachand, F. and Tian, B. (2017) Comparative analysis of alternative polyadenylation in S. Cerevisiae and S. Pombe. Genome Res., 27, 1685–1695.

64. Rodríguez-Romero, J., Marconi, M., Ortega-Campayo, V., Demuez, M., Wilkinson, M.D. and Sesma, A. (2019) Virulence- and signaling-associated genes display a preference for long 3′UTRs during rice infection and metabolic stress in the rice blast fungus. New Phytol., 221, 399–414.

65. Zhao, J., Hyman, L. and Moore, C. (1999) Formation of mRNA 3′ Ends in Eukaryotes: Mechanism, Regulation, and Interrelationships with Other Steps in mRNA Synthesis. Microbiol. Mol. Biol. Rev., 63, 405–445.

66. Neafsey, D.E. and Galagan, J.E. (2007) Dual modes of natural selection on upstream open reading frames. Mol. Biol. Evol., 24, 1744–1751.

67. Galagan, J.E., Calvo, S.E., Cuomo, C., Ma, L.-J., Wortman, J.R., Batzoglou, S., Lee, S.-I., Baştürkmen, M., Spevak, C.C., Clutterbuck, J., et al. (2005) Sequencing of Aspergillus nidulans and comparative analysis with A. fumigatus and A. oryzae. Nature, 438, 1105– 1115.

68. Dever, T.E., Ivanov, I.P. and Sachs, M.S. (2020) Conserved Upstream Open Reading Frame Nascent Peptides That Control Translation. Annu. Rev. Genet., 54, 237–264.

69. Spevak, C.C., Ivanov, I.P. and Sachs, M.S. (2010) Sequence requirements for ribosome stalling by the arginine attenuator peptide. J. Biol. Chem., 285, 40933–40942.

70. Sinturel, F., Navickas, A., Wery, M., Descrimes, M., Morillon, A., Torchet, C. and Benard, L. (2015) Cytoplasmic Control of Sense-Antisense mRNA Pairs. Cell Rep., 12, 1853– 1864.

71. Prescott, E.M. and Proudfoot, N.J. (2002) Transcriptional collision between convergent genes in budding yeast. Proc. Natl. Acad. Sci. U. S. A., 99, 8796–8801.

72. Fox-Walsh, K.L. and Hertel, K.J. (2009) Splice-site pairing is an intrinsically high fidelity process. Proc. Natl. Acad. Sci., 106, 1766–1771.

73. Grützmann, K., Szafranski, K., Pohl, M., Voigt, K., Petzold, A. and Schuster, S. (2014) Fungal alternative splicing is associated with multicellular complexity and virulence: A genome-wide multi-species study. DNA Res., 21, 27–39.

74. Xie, Y., Chan, P.-L., Kwan, H.-S. and Chang, J. (2023) The Genome-Wide Characterization of Alternative Splicing and RNA Editing in the Development of Coprinopsis cinerea. J. Fungi, 9, 915.

75. Fang, S., Hou, X., Qiu, K., He, R., Feng, X. and Liang, X. (2020) The occurrence and function of alternative splicing in fungi. Fungal Biol. Rev., 34, 178–188.

76. Corley, S.M., Troy, N.M., Bosco, A. and Wilkins, M.R. (2019) QuantSeq. 3’ Sequencing combined with Salmon provides a fast, reliable approach for high throughput RNA expression analysis. Sci. Rep., 9, 18895.

77. Xu, C. and Zhang, J. (2018) Alternative Polyadenylation of Mammalian Transcripts Is Generally Deleterious, Not Adaptive. Cell Syst., 6, 734–742.e4.

78. Nagy, L.G., Vonk, P.J., Künzler, M., Földi, C., Virágh, M., Ohm, R.A., Hennicke, F., Bálint, B., Csemetics, Á., Hegedüs, B., et al. (2023) Lessons on fruiting body morphogenesis from genomes and transcriptomes of Agaricomycetes. Stud. Mycol., 104, 1–85.

79. Pareek, M., Hegedüs, B., Hou, Z., Csernetics, Á., Wu, H., Virágh, M., Sahu, N., Liu, X.-B. and Nagy, L. (2022) Preassembled Cas9 Ribonucleoprotein-Mediated Gene Deletion Identifies the Carbon Catabolite Repressor and Its Target Genes in Coprinopsis cinerea. Appl. Environ. Microbiol., 10.1128/aem.00940-22.

80. Marian, I.M., Vonk, P.J., Valdes, I.D., Barry, K., Bostock, B., Carver, A., Daum, C., Lerner, H., Lipzen, A., Park, H., et al. (2022) The Transcription Factor Roc1 Is a Key Regulator of Cellulose Degradation in the Wood-Decaying Mushroom Schizophyllum commune. mBio, 13, e00628–22.

81. Cai, P., Wang, B., Ji, J., Jiang, Y., Wan, L., Tian, C. and Ma, Y. (2015) The Putative Cellodextrin Transporter-like Protein CLP1 Is Involved in Cellulase Induction in Neurospora crassa*. J. Biol. Chem., 290, 788–796.

82. Znameroski, E.A., Li, X., Tsai, J.C., Galazka, J.M., Glass, N.L. and Cate, J.H.D. (2014) Evidence for Transceptor Function of Cellodextrin Transporters in Neurospora crassa*. J. Biol. Chem., 289, 2610–2619.

83. Kuratani, M., Tanaka, K., Terashima, K., Muraguchi, H., Nakazawa, T., Nakahori, K. and Kamada, T. (2010) The dst2 gene essential for photomorphogenesis of Coprinopsis cinerea encodes a protein with a putative FAD-binding-4 domain. Fungal Genet. Biol., 47, 152–158.

84. Boulianne, R.P., Liu, Y., Aebi, M., Lu, B.C. and Kues, U. (2000) Fruiting body development in Coprinus cinereus: Regulated expression of two galectins secreted by a non-classical pathway. Microbiology, 146, 1841–1853.

85. Sakamoto, Y., Sato, S., Ito, M., Ando, Y., Nakahori, K. and Muraguchi, H. (2018) Blue light exposure and nutrient conditions influence the expression of genes involved in simultaneous hyphal knot formation in Coprinopsis cinerea. Microbiol. Res., 217, 81– 90.

86. van Munster, J.M., Daly, P., Delmas, S., Pullan, S.T., Blythe, M.J., Malla, S., Kokolski, M., Noltorp, E.C.M., Wennberg, K., Fetherston, R., et al. (2014) The role of carbon starvation in the induction of enzymes that degrade plant-derived carbohydrates in Aspergillus niger. Fungal Genet. Biol., 72, 34–47.

87. Ellström, M., Shah, F., Johansson, T., Ahrén, D., Persson, P. and Tunlid, A. (2015) The carbon starvation response of the ectomycorrhizal fungus Paxillus involutus. FEMS Microbiol. Ecol., 91, fiv027.

88. Nitsche, B.M., Jørgensen, T.R., Akeroyd, M., Meyer, V. and Ram, A.F.J. (2012) The carbon starvation response of Aspergillus niger during submerged cultivation: Insights from the transcriptome and secretome. BMC Genomics, 13.

89. Glass, N.L., Schmoll, M., Cate, J.H.D. and Coradetti, S. (2013) Plant Cell Wall Deconstruction by Ascomycete Fungi. Annu. Rev. Microbiol., 67, 477–498.

90. Benz, J.P., Chau, B.H., Zheng, D., Bauer, S., Glass, N.L. and Somerville, C.R. (2014) A comparative systems analysis of polysaccharide-elicited responses in Neurospora crassa reveals carbon source-specific cellular adaptations. Mol. Microbiol., 91, 275–299.

91. Coradetti, S.T., Craig, J.P., Xiong, Y., Shock, T., Tian, C. and Glass, N.L. (2012) Conserved and essential transcription factors for cellulase gene expression in ascomycete fungi. Proc. Natl. Acad. Sci., 109, 7397–7402.

92. Delmas, S., Pullan, S.T., Gaddipati, S., Kokolski, M., Malla, S., Blythe, M.J., Ibbett, R., Campbell, M., Liddell, S., Aboobaker, A., et al. (2012) Uncovering the Genome-Wide Transcriptional Responses of the Filamentous Fungus Aspergillus niger to Lignocellulose Using RNA Sequencing. PLOS Genet., 8, e1002875.

93. Ando, Y., Nakazawa, T., Oka, K., Nakahori, K. and Kamada, T. (2013) Cc.snf5, a gene encoding a putative component of the SWI/SNF chromatin remodeling complex, is essential for sexual development in the agaricomycete Coprinopsis cinerea. Fungal Genet. Biol., 50, 82–89.

94. Liu, C., Kang, L., Lin, M., Bi, J., Liu, Z. and Yuan, S. (2022) Molecular Mechanism by Which the GATA Transcription Factor CcNsdD2 Regulates the Developmental Fate of Coprinopsis cinerea under Dark or Light Conditions. mBio, 13.

95. Nakazawa, T., Ando, Y., Kitaaki, K., Nakahori, K. and Kamada, T. (2011) Efficient gene targeting in ΔCc.ku70 or ΔCc.lig4 mutants of the agaricomycete Coprinopsis cinerea. Fungal Genet. Biol., 48, 939–946.

96. Brych, A., Mascarenhas, J., Jaeger, E., Charkiewicz, E., Pokorny, R., Bölker, M., Doehlemann, G. and Batschauer, A. (2016) White collar 1-induced photolyase expression contributes to UV-tolerance of Ustilago maydis. MicrobiologyOpen, 5, 224– 243.

97. Ohm, R.A., Aerts, D., Wösten, H.A.B. and Lugones, L.G. (2013) The blue light receptor complex WC-1/2 of Schizophyllum commune is involved in mushroom formation and protection against phototoxicity. Environ. Microbiol., 15, 943–955.

98. Froehlich, A.C., Chen, C.H., Belden, W.J., Madeti, C., Roenneberg, T., Merrow, M., Loros, J.J. and Dunlap, J.C. (2010) Genetic and molecular characterization of a cryptochrome from the filamentous fungus Neurospora crassa. Eukaryot. Cell, 9, 738–750.

99. Chen, C., Ringelberg, C.S., Gross, R.H., Dunlap, J.C. and Loros, J.J. (2009) Genome-wide analysis of light-inducible responses reveals hierarchical light signalling in Neurospora. EMBO J., 28, 1029–1042.

100. Wu, C., Yang, F., Smith, K.M., Peterson, M., Dekhang, R., Zhang, Y., Zucker, J., Bredeweg, E.L., Mallappa, C., Zhou, X., et al. (2014) Genome-Wide Characterization of Light-Regulated Genes in Neurospora crassa. G3 GenesGenomesGenetics, 4, 1731–1745.

101. Smith, K.M., Sancar, G., Dekhang, R., Sullivan, C.M., Li, S., Tag, A.G., Sancar, C., Bredeweg, E.L., Priest, H.D., McCormick, R.F., et al. (2010) Transcription factors in light and circadian clock signaling networks revealed by genomewide mapping of direct targets for neurospora white collar complex. Eukaryot. Cell, 9, 1549–1556.

102. Sano, H., Kaneko, S., Sakamoto, Y., Sato, T. and Shishido, K. (2009) The basidiomycetous mushroom Lentinula edodes white collar-2 homolog PHRB, a partner of putative blue-light photoreceptor PHRA, binds to a specific site in the promoter region of the L. edodes tyrosinase gene. Fungal Genet. Biol., 46, 333–341.

103. Pelkmans, J.F., Lugones, L.G. and Wösten, H.A.B. (2016) 15 Fruiting Body Formation in Basidiomycetes. In Wendland, J. (ed), Growth, Differentiation and Sexuality, The Mycota. Springer International Publishing, Cham, pp. 387–405.

104. Wösten, H.A.B., Wetter, M.-A. van, Lugones, L.G., Mei, H.C. van der, Busscher, H.J. and Wessels, J.G.H. (1999) How a fungus escapes the water to grow into the air. Curr. Biol., 9, 85–88.

105. Erdmann, S., Freihorst, D., Raudaskoski, M., Schmidt-Heck, W., Jung, E.-M., Senftleben, D. and Kothe, E. (2012) Transcriptome and Functional Analysis of Mating in the Basidiomycete Schizophyllum commune. Eukaryot. Cell, 11, 571–589.

106. Sammer, D., Krause, K., Gube, M., Wagner, K. and Kothe, E. (2016) Hydrophobins in the Life Cycle of the Ectomycorrhizal Basidiomycete Tricholoma vaccinum. PLOS ONE, 11, e0167773.

107. Bayry, J., Aimanianda, V., Guijarro, J.I., Sunde, M. and Latgé, J.-P. (2012) Hydrophobins— Unique Fungal Proteins. PLoS Pathog., 8, e1002700.

108. Ohm, R.A., de Jong, J.F., de Bekker, C., Wösten, H.A.B. and Lugones, L.G. (2011) Transcription factor genes of Schizophyllum commune involved in regulation of mushroom formation. Mol. Microbiol., 81, 1433–1445.

109. Pelkmans, J.F., Patil, M.B., Gehrmann, T., Reinders, M.J.T., Wösten, H.A.B. and Lugones, L.G. (2017) Transcription factors of schizophyllum commune involved in mushroom formation and modulation of vegetative growth. Sci. Rep., 7, 1–11.

110. Plaza, D.F., Lin, C.W., van der Velden, N.S.J., Aebi, M. and Künzler, M. (2014) Comparative transcriptomics of the model mushroom Coprinopsis cinerea reveals tissue-specific armories and a conserved circuitry for sexual development. BMC Genomics, 15, 1–17.

111. Kombrink, A., Tayyrov, A., Essig, A., Stöckli, M., Micheller, S., Hintze, J., van Heuvel, Y., Dürig, N., Lin, C. wei, Kallio, P.T., et al. (2019) Induction of antibacterial proteins and peptides in the coprophilous mushroom Coprinopsis cinerea in response to bacteria. ISME J., 13, 588–602.

112. Xie, Y., Chang, J. and Kwan, H.S. (2020) Carbon metabolism and transcriptome in developmental paths differentiation of a homokaryotic Coprinopsis cinerea strain. Fungal Genet. Biol., 143, 103432.

113. Salzberg, S.L. (2019) Next-generation genome annotation: we still struggle to get it right. Genome Biol., 20, 92.

114. Chen, C.-H. and Loros, J.J. (2009) Neurospora sees the light: Light signaling components in a model system. Commun. Integr. Biol., 2, 448–451.

115. Kodzius, R., Kojima, M., Nishiyori, H., Nakamura, M., Fukuda, S., Tagami, M., Sasaki, D., Imamura, K., Kai, C., Harbers, M., et al. (2006) CAGE: cap analysis of gene expression. Nat. Methods, 3, 211–222.

116. Földi, C., Merényi, Z., Balázs, B., Csernetics, Á., Miklovics, N., Wu, H., Hegedüs, B., Virágh, M., Hou, Z., Liu, X.-B., et al. (2023) Snowball: a novel gene family required for developmental patterning in fruiting bodies of mushroom-forming fungi (Agaricomycetes).10.1101/2023.11.13.566867.

117. Swamy, S., Uno, I. and Ishikawa, T. (1984) Morphogenetic Effects of Mutations at the A and B Incompatibility Factors in Coprinus cinereus. Microbiology, 130, 3219–3224.

118. Vaser, R., Sović, I., Nagarajan, N. and Šikić, M. (2017) Fast and accurate de novo genome assembly from long uncorrected reads. Genome Res., 27, 737–746.

119. Kolmogorov, M., Yuan, J., Lin, Y. and Pevzner, P.A. (2019) Assembly of long, error-prone reads using repeat graphs. Nat. Biotechnol., 37, 540–546.

120. Bushnell, B., Rood, J. and Singer, E. (2017) BBMerge – Accurate paired shotgun read merging via overlap. PLOS ONE, 12, e0185056.

121. Bengtsson-Palme, J., Ryberg, M., Hartmann, M., Branco, S., Wang, Z., Godhe, A., De Wit, P., Sánchez-García, M., Ebersberger, I., de Sousa, F., et al. (2013) Improved software detection and extraction of ITS1 and ITS2 from ribosomal ITS sequences of fungi and other eukaryotes for analysis of environmental sequencing data. Methods Ecol. Evol., 4, 914–919.

122. epi2me-labs/pychopper (2024).

123. Salmela, L. and Rivals, E. (2014) LoRDEC: Accurate and efficient long read error correction. Bioinformatics, 30, 3506–3514.

124. Durai, D.A. and Schulz, M.H. (2019) Improving in-silico normalization using read weights. Sci. Rep., 9, 1–10.

125. nanoporetech/pomoxis (2024).

126. Marçais, G., Delcher, A.L., Phillippy, A.M., Coston, R., Salzberg, S.L. and Zimin, A. (2018) MUMmer4: A fast and versatile genome alignment system. PLoS Comput. Biol., 14, 1– 14.

127. Goel, M., Sun, H., Jiao, W.B. and Schneeberger, K. (2019) SyRI: finding genomic rearrangements and local sequence differences from whole-genome assemblies. Genome Biol., 20, 1–13.

128. Goel, M. and Schneeberger, K. (2022) Plotsr: Visualizing Structural Similarities and Rearrangements Between Multiple Genomes. Bioinformatics, 38, 2922–2926.

129. Chen, S., Zhou, Y., Chen, Y. and Gu, J. (2018) fastp: an ultra-fast all-in-one FASTQ preprocessor. Bioinformatics, 34, i884–i890.

130. Li, H. and Durbin, R. (2009) Fast and accurate short read alignment with Burrows-Wheeler transform. Bioinforma. Oxf. Engl., 25, 1754–1760.

131. Danecek, P., Bonfield, J.K., Liddle, J., Marshall, J., Ohan, V., Pollard, M.O., Whitwham, A., Keane, T., McCarthy, S.A., Davies, R.M., et al. (2021) Twelve years of SAMtools and BCFtools. GigaScience, 10, giab008.

132. broadinstitute/picard (2024).

133. McKenna, A., Hanna, M., Banks, E., Sivachenko, A., Cibulskis, K., Kernytsky, A., Garimella, K., Altshuler, D., Gabriel, S., Daly, M., et al. (2010) The Genome Analysis Toolkit: A MapReduce framework for analyzing next-generation DNA sequencing data. Genome Res., 20, 1297–1303.

134. Liu, B., Liu, Y., Li, J., Guo, H., Zang, T. and Wang, Y. (2019) DeSALT: Fast and accurate long transcriptomic read alignment with De Bruijn graph-based index. Genome Biol., 20, 1–14.

135. Li, H. Minimap2: pairwise alignment for nucleotide sequences. 10.1093/bioinformatics/bty191.

136. Boratyn, G.M., Thierry-Mieg, J., Thierry-Mieg, D., Busby, B. and Madden, T.L. (2019) Magic-BLAST, an accurate RNA-seq aligner for long and short reads. BMC Bioinformatics, 20, 1–19.

137. Marić, J., Sović, I., Križanović, K., Nagarajan, N. and Šikić, M. (2019) Graphmap2 - splice-aware RNA-seq mapper for long reads. bioRxiv, 10.1101/720458.

138. Kuo, R.I., Cheng, Y., Zhang, R., Brown, J.W.S., Smith, J., Archibald, A.L. and Burt, D.W. (2020) Illuminating the dark side of the human transcriptome with long read transcript sequencing. BMC Genomics, 21, 1–22.

139. Tjeldnes, H., Labun, K., Torres Cleuren, Y., Chyżyńska, K., Świrski, M. and Valen, E. (2021) ORFik: a comprehensive R toolkit for the analysis of translation. BMC Bioinformatics, 22, 336.

140. Tardaguila, M., Fuente, L. de la, Marti, C., Pereira, C., Pardo-Palacios, F.J., Risco, H. del, Ferrell, M., Mellado, M., Macchietto, M., Verheggen, K., et al. (2018) SQANTI: extensive characterization of long-read transcript sequences for quality control in full-length transcriptome identification and quantification. Genome Res., 28, 396–411.

141. Waterhouse, R.M., Seppey, M., Simão, F.A., Manni, M., Ioannidis, P., Klioutchnikov, G., Kriventseva, E.V. and Zdobnov, E.M. (2018) BUSCO Applications from Quality Assessments to Gene Prediction and Phylogenomics. Mol. Biol. Evol., 35, 543–548.

142. Trincado, J.L., Entizne, J.C., Hysenaj, G., Singh, B., Skalic, M., Elliott, D.J. and Eyras, E. (2018) SUPPA2: Fast, accurate, and uncertainty-aware differential splicing analysis across multiple conditions. Genome Biol., 19, 1–11.

143. Wagih, O. (2017) ggseqlogo: a versatile R package for drawing sequence logos. Bioinformatics, 33, 3645–3647.

144. Signal, B., Gloss, B.S., Dinger, M.E. and Mercer, T.R. (2018) Machine learning annotation of human branchpoints. Bioinformatics, 34, 920–927.

145. Routh, A. (2019) DPAC: A tool for differential poly(A)-cluster usage from poly(A)- targeted RNAseq data. G3 Genes Genomes Genet., 9, 1825–1830.

146. Oksanen, J., Simpson, G.L., Blanchet, F.G., Kindt, R., Legendre, P., Minchin, P.R., O’Hara, R.B., Solymos, P., Stevens, M.H.H., Szoecs, E., et al. (2022) vegan: Community Ecology Package.

147. Wickham, H., Chang, W., Henry, L., Pedersen, T.L., Takahashi, K., Wilke, C., Woo, K., Yutani, H., Dunnington, D., Posit, et al. (2023) ggplot2: Create Elegant Data Visualisations Using the Grammar of Graphics.

148. Wood, S.N. (2017) Generalized Additive Models: An Introduction with R, Second Edition 2nd ed. Chapman and Hall/CRC, Boca Raton.

149. Andrews, S. (2010) FastQC A Quality Control tool for High Throughput Sequence Data.

150. Ewels, P., Magnusson, M., Lundin, S. and Käller, M. (2016) MultiQC: summarize analysis results for multiple tools and samples in a single report. Bioinformatics, 32, 3047–3048.

151. Dobin, A., Davis, C.A., Schlesinger, F., Drenkow, J., Zaleski, C., Jha, S., Batut, P., Chaisson, M. and Gingeras, T.R. (2013) STAR: Ultrafast universal RNA-seq aligner. Bioinformatics, 29, 15–21.

152. Liao, Y., Smyth, G.K. and Shi, W. (2014) featureCounts: an efficient general purpose program for assigning sequence reads to genomic features. Bioinformatics, 30, 923– 930.

153. Robinson, M.D., McCarthy, D.J. and Smyth, G.K. (2010) edgeR: a Bioconductor package for differential expression analysis of digital gene expression data. Bioinformatics, 26, 139–140.

154. Ritchie, M.E., Phipson, B., Wu, D., Hu, Y., Law, C.W., Shi, W. and Smyth, G.K. (2015) limma powers differential expression analyses for RNA-sequencing and microarray studies. Nucleic Acids Res., 43, e47.

155. Gu, Z. (2022) Complex heatmap visualization. iMeta, 1, e43.

156. Jones, P., Binns, D., Chang, H.-Y., Fraser, M., Li, W., McAnulla, C., McWilliam, H., Maslen, J., Mitchell, A., Nuka, G., et al. (2014) InterProScan 5: genome-scale protein function classification. Bioinformatics, 30, 1236–1240.

157. Zheng, J., Ge, Q., Yan, Y., Zhang, X., Huang, L. and Yin, Y. (2023) dbCAN3: automated carbohydrate-active enzyme and substrate annotation. Nucleic Acids Res., 51, W115– W121.

158. Sahu, N., Indic, B., Wong-Bajracharya, J., Merényi, Z., Ke, H.-M., Ahrendt, S., Monk, T.-L., Kocsubé, S., Drula, E., Lipzen, A., et al. (2023) Vertical and horizontal gene transfer shaped plant colonization and biomass degradation in the fungal genus Armillaria. Nat. Microbiol., 8, 1668–1681.

159. topGO Bioconductor.

160. GO.db Bioconductor.

