## Supplementary Figures for "Unraveling Morphogenesis, Starvation, and Light Responses in a Mushroom-Forming Fungus, *Coprinopsis cinerea*, Using Long Read Sequencing and Extensive Expression Profiling"

### Supplementary data

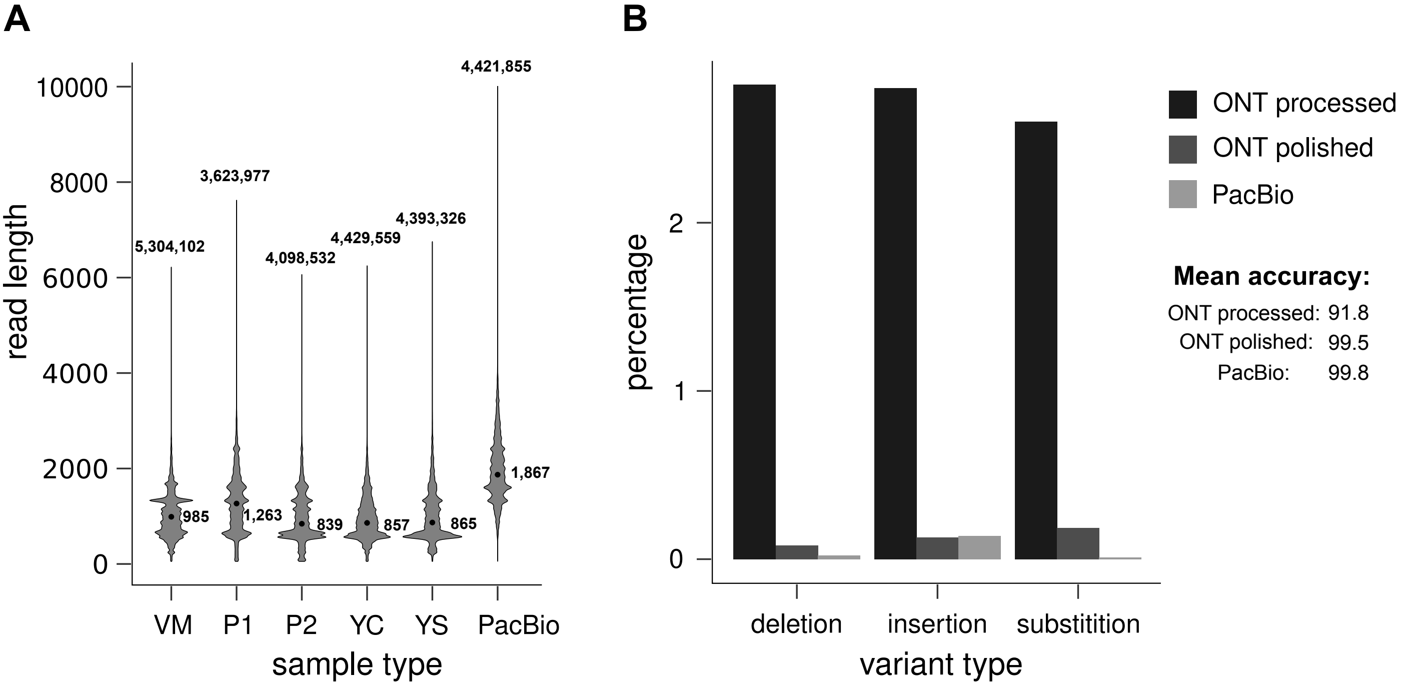


**Supplementary Figure 1.**
**Quantitative and qualitative comparison of the long-read libraries**
**A. Long-read cDNA library read length and read number distribution.** Median read lengths are marked with dots. Abbreviations of developmental stages are as follows for the ONT samples: VM -vegetative mycelium, P1 - stage 1 primordium, P2 - stage 2 primordium, YC - young fruiting body cap, YS - young fruiting body stipe.
**B. Assessment of the insertion, deletion and substitution rate of the long-read cDNA libraries by comparison to the CopciAB V2 genome using pomoxis.**

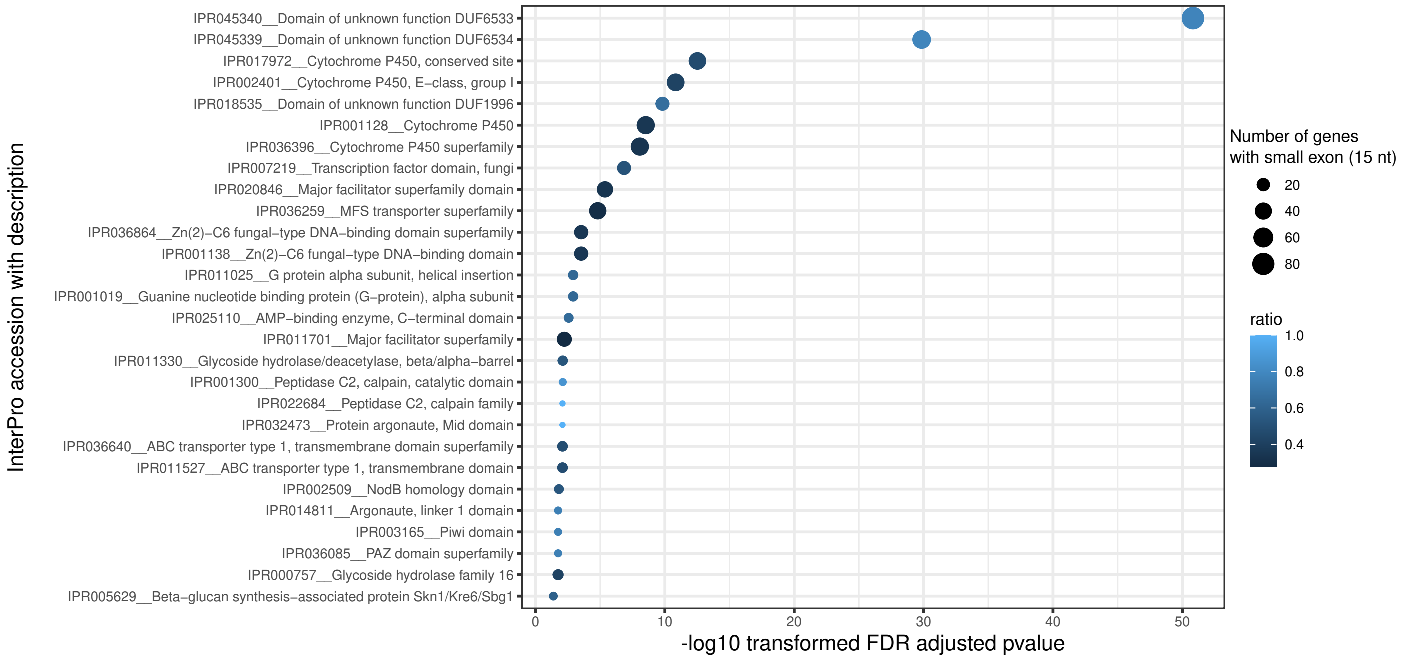


**Supplementary Figure 2.**

**Microexon enrichment between different InterPro families.** The y-axis represents the InterPro families that show significant enrichment (determined by Fisher exact test with FDR-adjusted *p* value ≤ 0.05) of microexons. The size of the dots reflects the number of genes within each family that contain a microexon. The shading of the dots indicates the ratio of microexon-containing genes within each family.

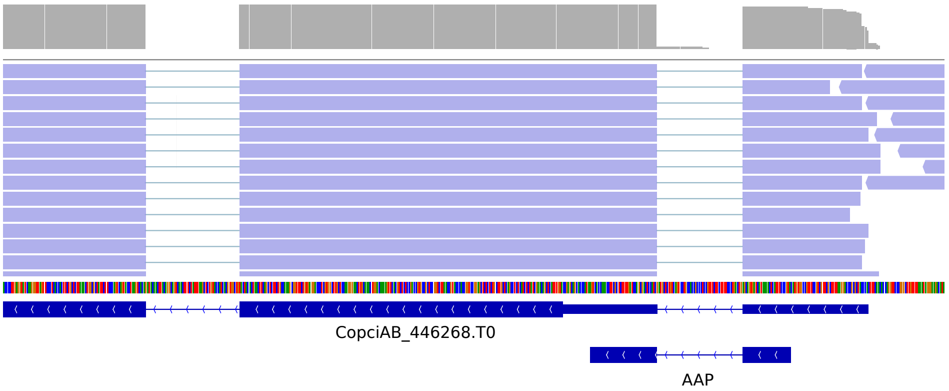


**Supplementary Figure 3.**

**5’ portion of the gene (CopciAB_446268.T0) orthologous to N. crassa arg-2 showing the putative arginine attenuator peptide (AAP) encoding uORF.** The figure was generated using IGV and illustrates the following features from top to bottom: the gray stripe represents the long-read coverage; the blue stripe below indicates individually aligned long-reads; the colored stripe below represents the nucleotide sequence of the genome; the blue boxes below the sequence correspond to the predicted gene models and uORF.

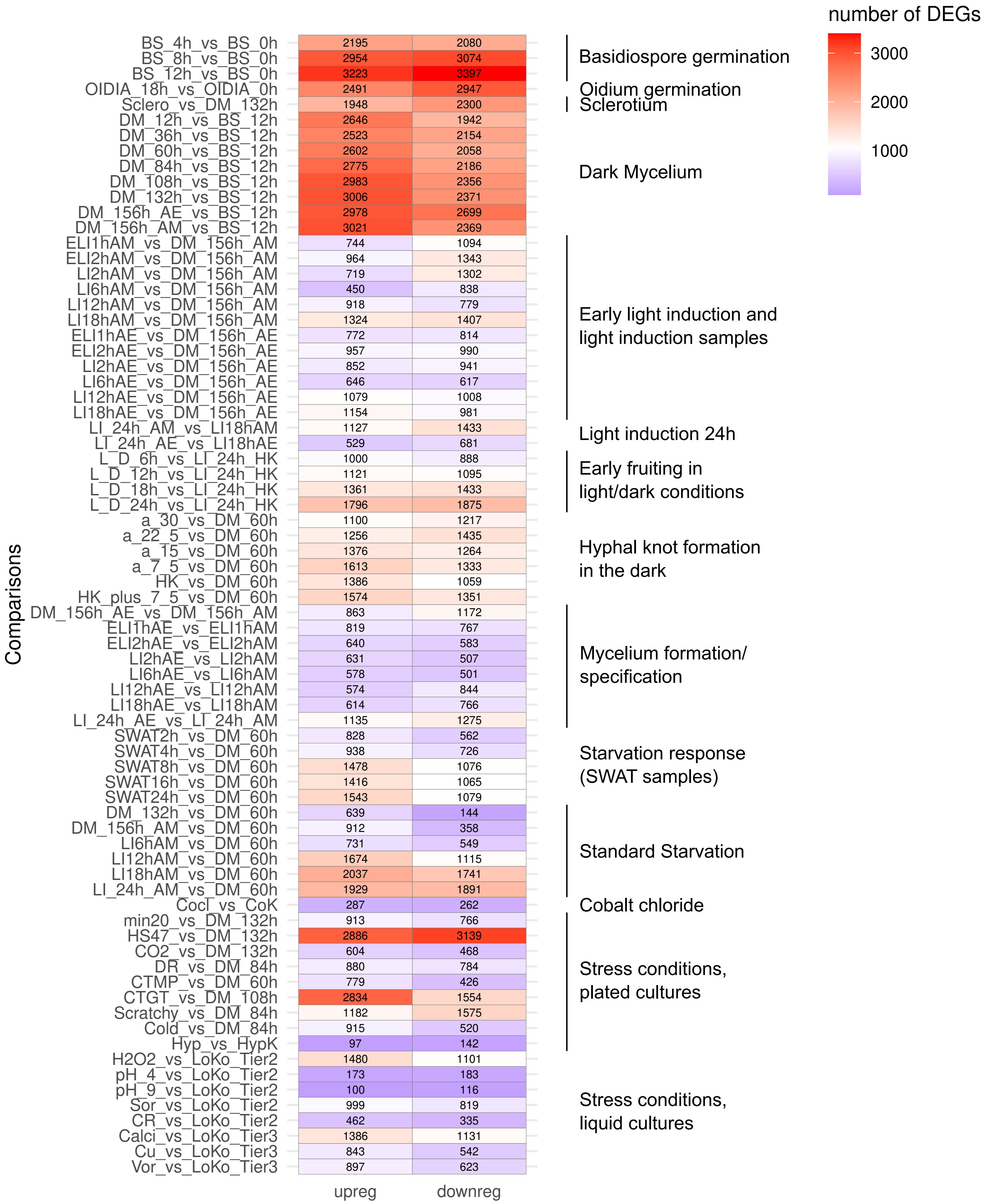


**Supplementary Figure 4.**
**Number of significantly (BH adjusted *p* ≤0.05, fold-change ≥2) differentially expressed genes (DEGs) obtained during the comparisons of RNA-Seq data.**

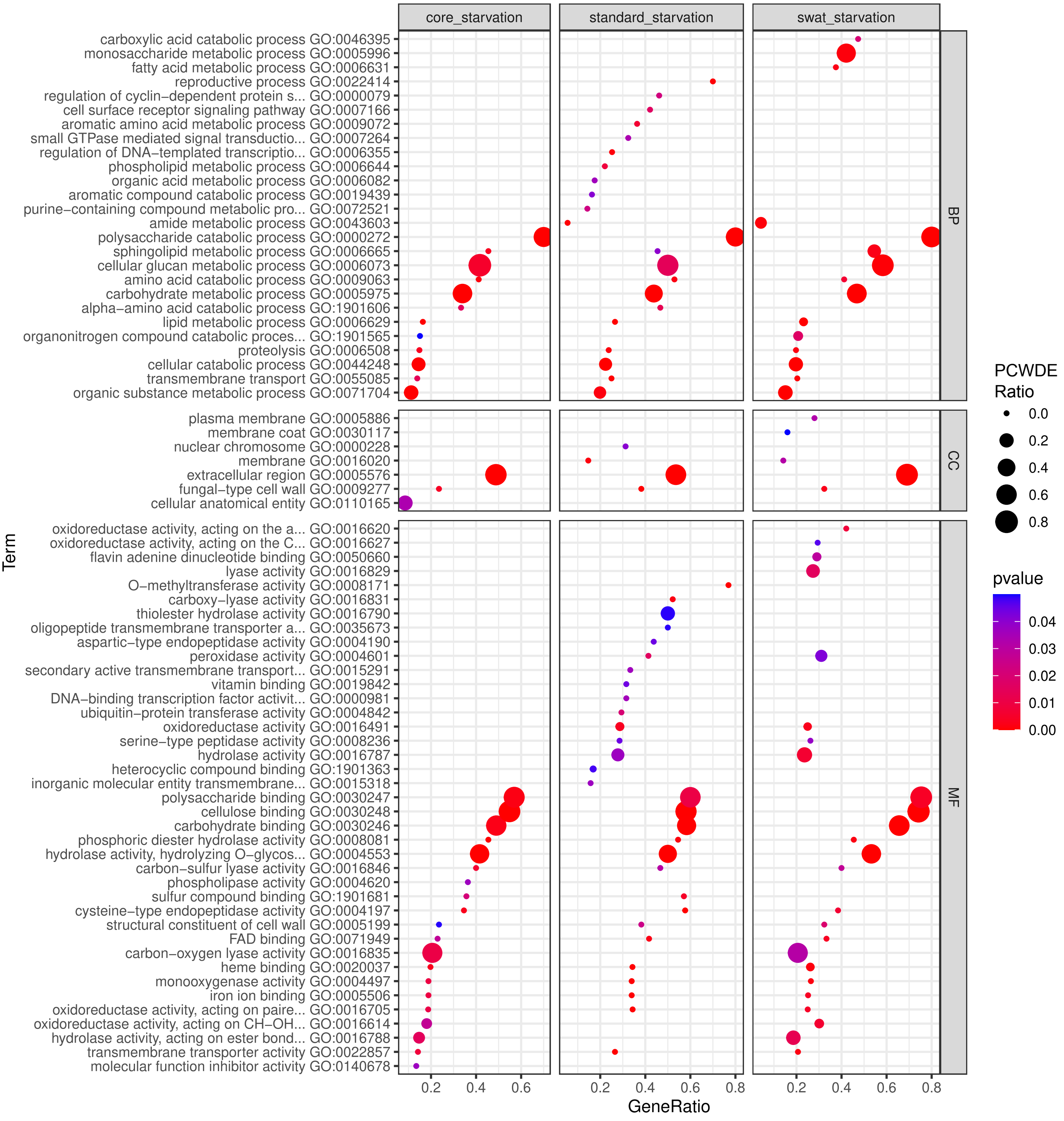


**Supplementary Figure 5.**
**Significantly enriched GO groups for core starvation, standard starvation, and SWAT starvation gene sets and the PCWDEs ratios within these groups.** The y-axis represents the ratio of GO groups present in the gene sets. The size of the dots corresponds to the PCWDE ratio of the GO groups present. The color of the dots indicates the statistical significance of the enrichment.

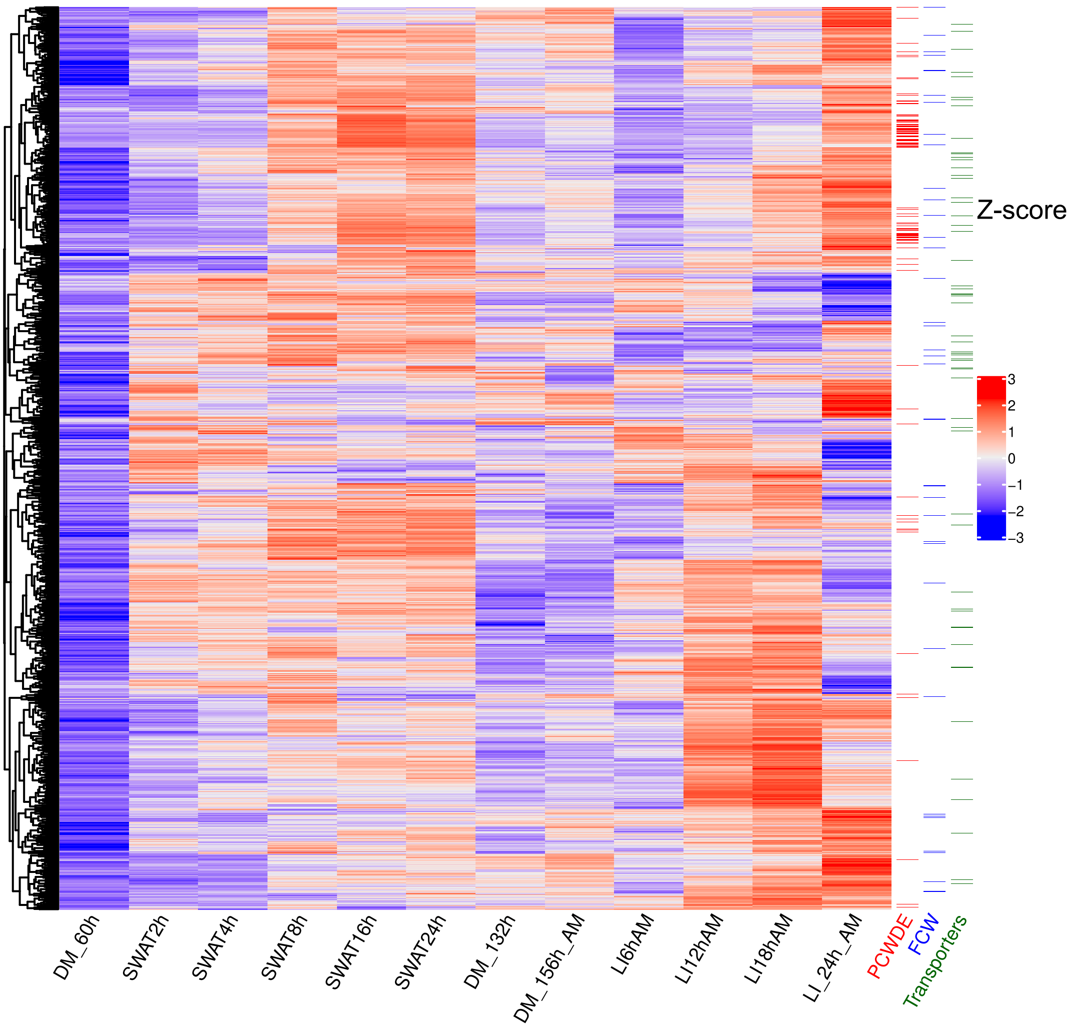


**Supplementary Figure 6.**
**Heatmap representation of the core starvation response gene expression in the standard and SWAT samples.** Gene expression values are represented by the mean $log_2$CPM of the biological replicates, which were scaled (Z-score) and hierarchically clustered by row (WardD method based on the Euclidean distance). 'DM', 'SWAT' and 'LI' stands for dark grown mycelium, Water agar transfer and Light induction, respectively the length of incubation time in the given condition in hours. In the last three columns of the figure, the genes associated with functions within the PCWDE, FCW, and Transports categories are highlighted.

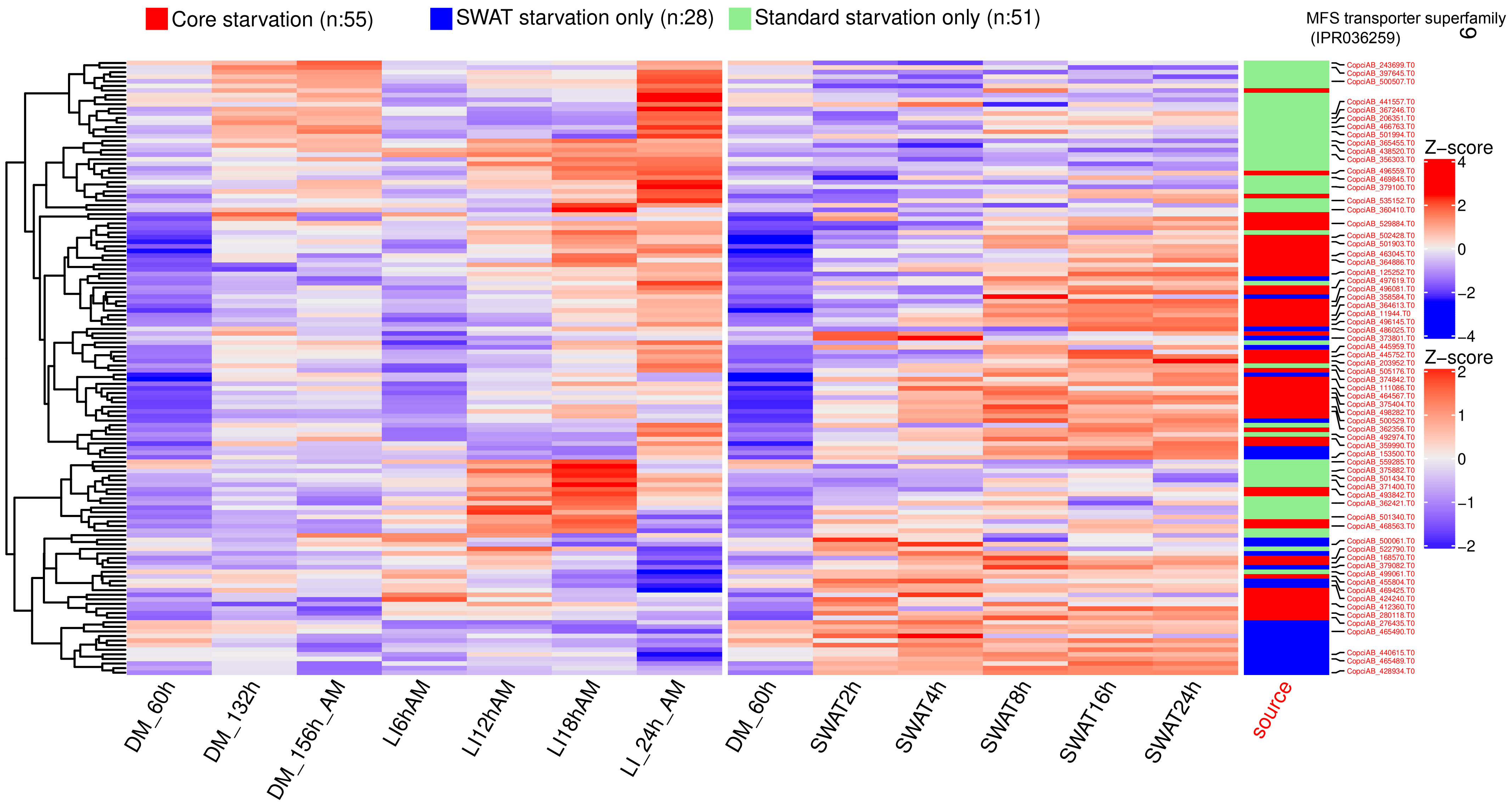


**Supplementary Figure 7.**
**Heatmap representation of the starvation-induced expression of putative transporter-encoding genes (n: 134) in the standard and SWAT samples.** Transporters were identified based on their membership in the GO:0055085 gene ontology group, supplemented by some additional manually curated proteins containing transmembrane domains. Gene expression values are represented by the mean $log_2$CPM of the biological replicants, which were scaled (Z-score) and hierarchically clustered by row (WardD method based on the Euclidean distance). In the last column, the transporters are highlighted according to their affiliation to the subcategories “Core starvation”, “SWAT starvation only” and “Standard starvation only”. Transporters belonging to the MFS supperfamily (IPR036259) are also highlighted. In the case of sample names, 'DM', 'SWAT' and 'LI' stands for dark grown mycelium, Water agar transfer and Light induction, respectively the length of incubation time in the given condition in hours. 'AM' stands for the attached mycelium.

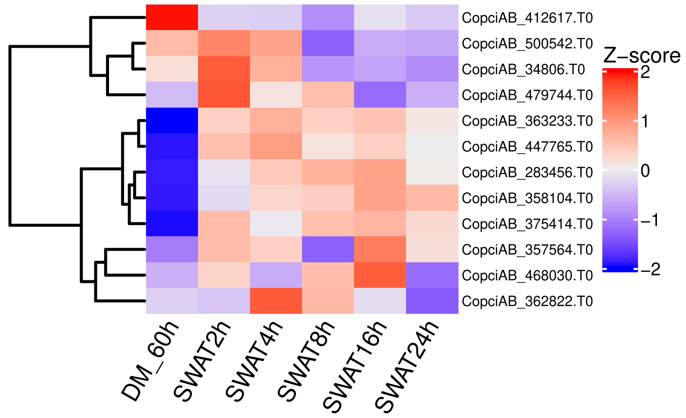


**Supplementary Figure 8.**

**Heatmap representation of the expression of the 12 putative orthologs (reciprocal best hits) of 35 previously reported autophagy genes of S. cerevisiae in the SWAT samples.** Gene expression values are represented by the mean $log_2$CPM of the biological replicates, which were scaled (Z-score) and hierarchically clustered by row (WardD method based on the Euclidean distance). In the case of sample names, 'DM' and 'SWAT' stands for dark grown mycelium and Water agar transfer, respectively the length of incubation time in the given condition in hours.

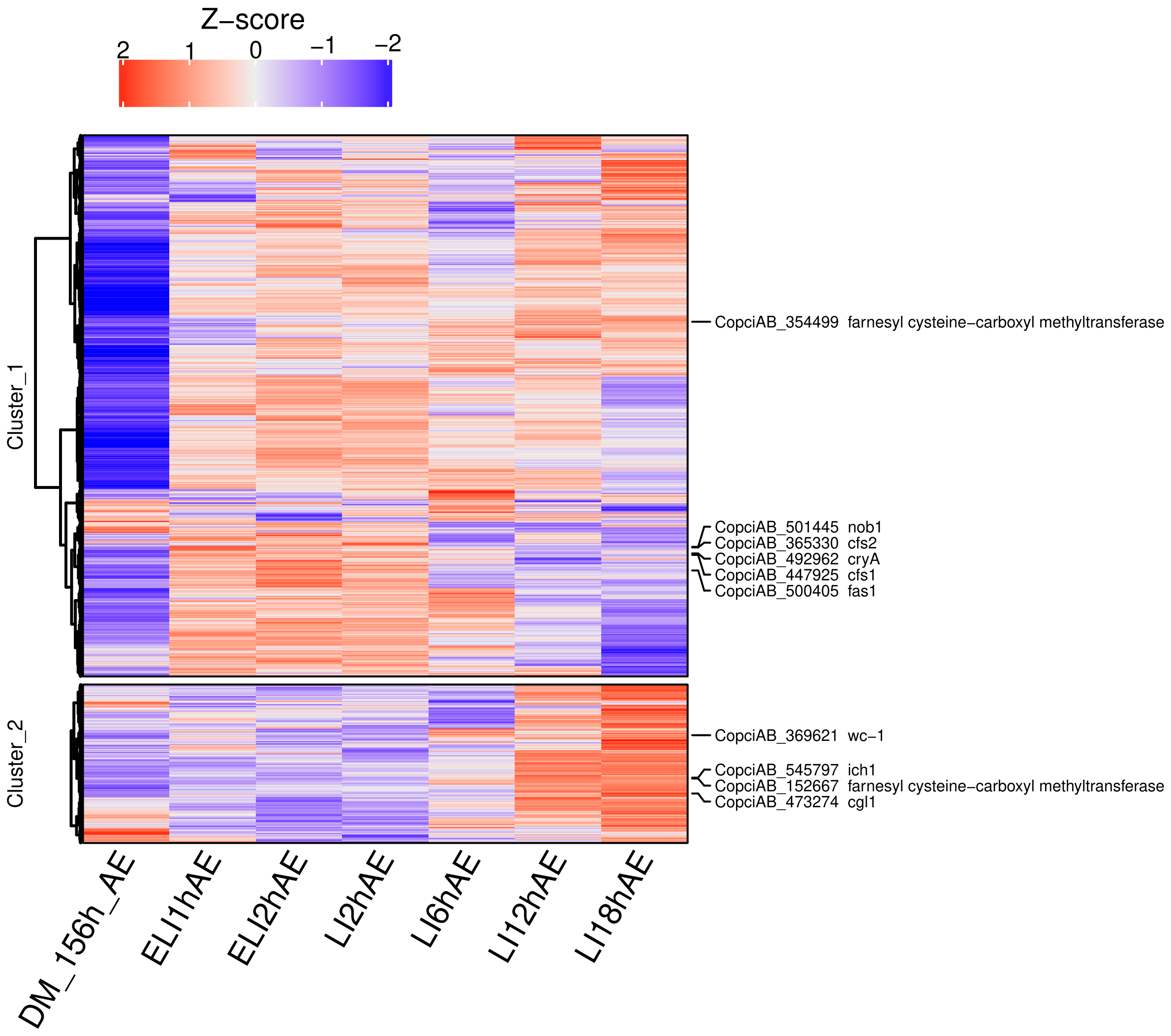


**Supplementary Figure 9.**

**Heatmap representation of the $log_2$CPM expression values of the core light response genes in the AE samples.** Genes were hierarchically clustered into two groups using the WardD method based on the Euclidean distance between the Z-score-normalized $log_2$CPM gene expression values. Cluster 1 captures the early light-induced genes, while cluster 2 captures the late light-induced genes. Genes identified in the literature as light-induced genes are marked on the right side of the figure. In the case of sample names, 'DM', 'ELI' and 'LI' stands for dark grown mycelium, Early Light induction and Light induction, respectively the length of incubation time in the given condition in hours. 'AE' stands for the aerial mycelium.

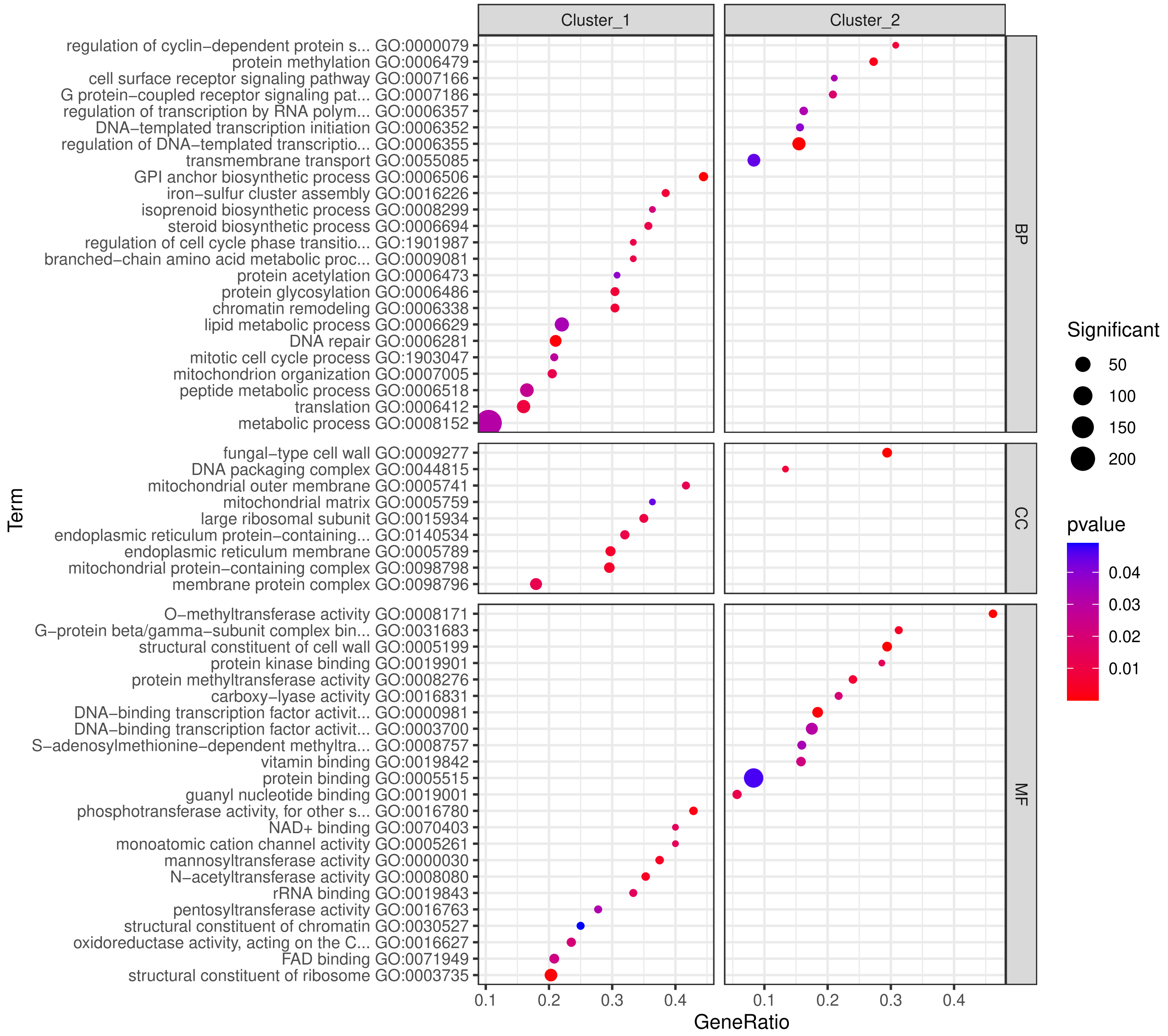


**Supplementary Figure 10.**
**Significantly enriched GO groups for core light response gene sets in the AM samples.** The y-axis represents the ratio of GO groups present in the gene sets. The size of the dots corresponds to the number of significantly expressed genes belonging to the GO group. The color of the dots indicates the statistical significance of the enrichment. The vertical segmentation separates the early (cluster 1) and late light-induced genes (cluster 2). The horizontal segmentation is based on the three main aspects of the Gene Ontology (GO) classification: Biological Process (BP), Cellular Component (CC), Molecular Function (MF).

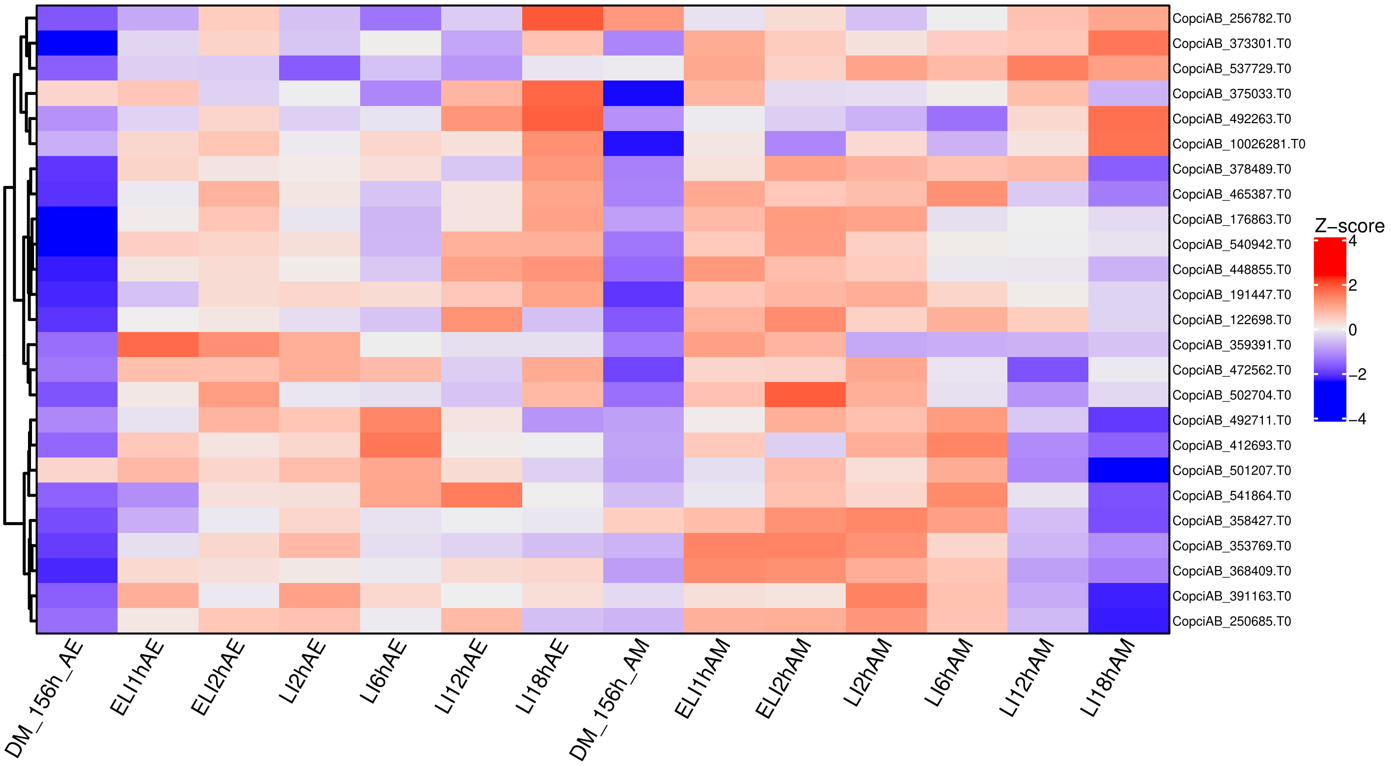


**Supplementary Figure 11.**
**Heatmap representation of the expression of the genes related to DNA-repair (GO:0006281) in the light induced samples.** Gene expression values are represented by the mean $\log_2$CPM of the biological replicants, which were scaled (Z-score) and hierarchically clustered by row (WardD method based on the Euclidean distance). In the case of sample names, 'DM', 'ELI' and 'LI' stands for dark grown mycelium, Early Light induction and Light induction, respectively the length of incubation time in the given condition in hours. 'AE' and 'AM' stands for the aerial mycelium and attached mycelium, respectively.

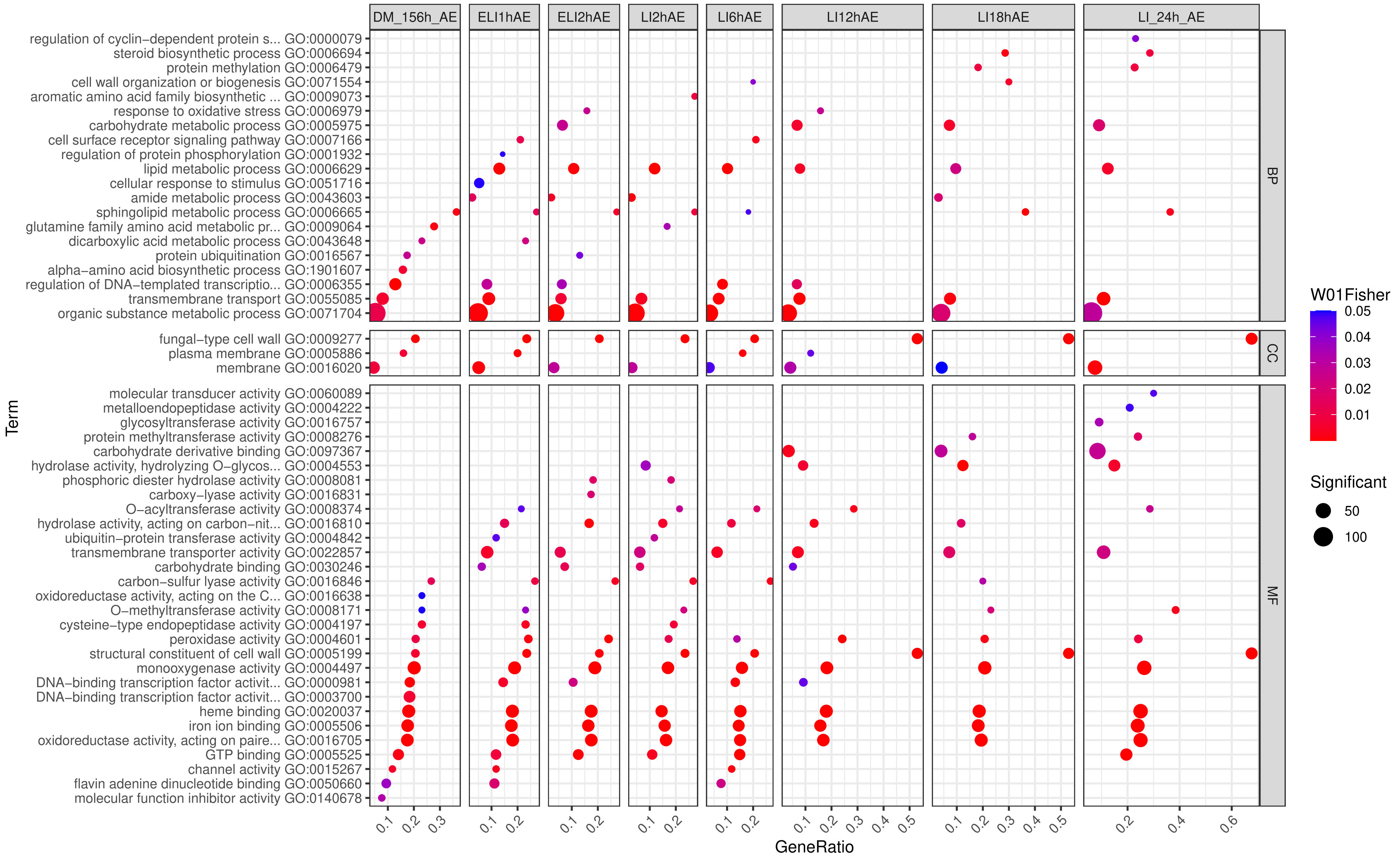


**Supplementary Figure 12.**

**Significantly enriched GO groups for light-induced aerial mycelium (AE) samples.** The y-axis represents the ratio of GO groups present in the gene sets. The size of the dots corresponds to the number of significantly expressed genes belonging to the GO group. The color of the dots indicates the statistical significance of the enrichment. The horizontal segmentation is based on the three main aspects of the Gene Ontology (GO) classification: Biological Process (BP), Cellular Component (CC), Molecular Function (MF).

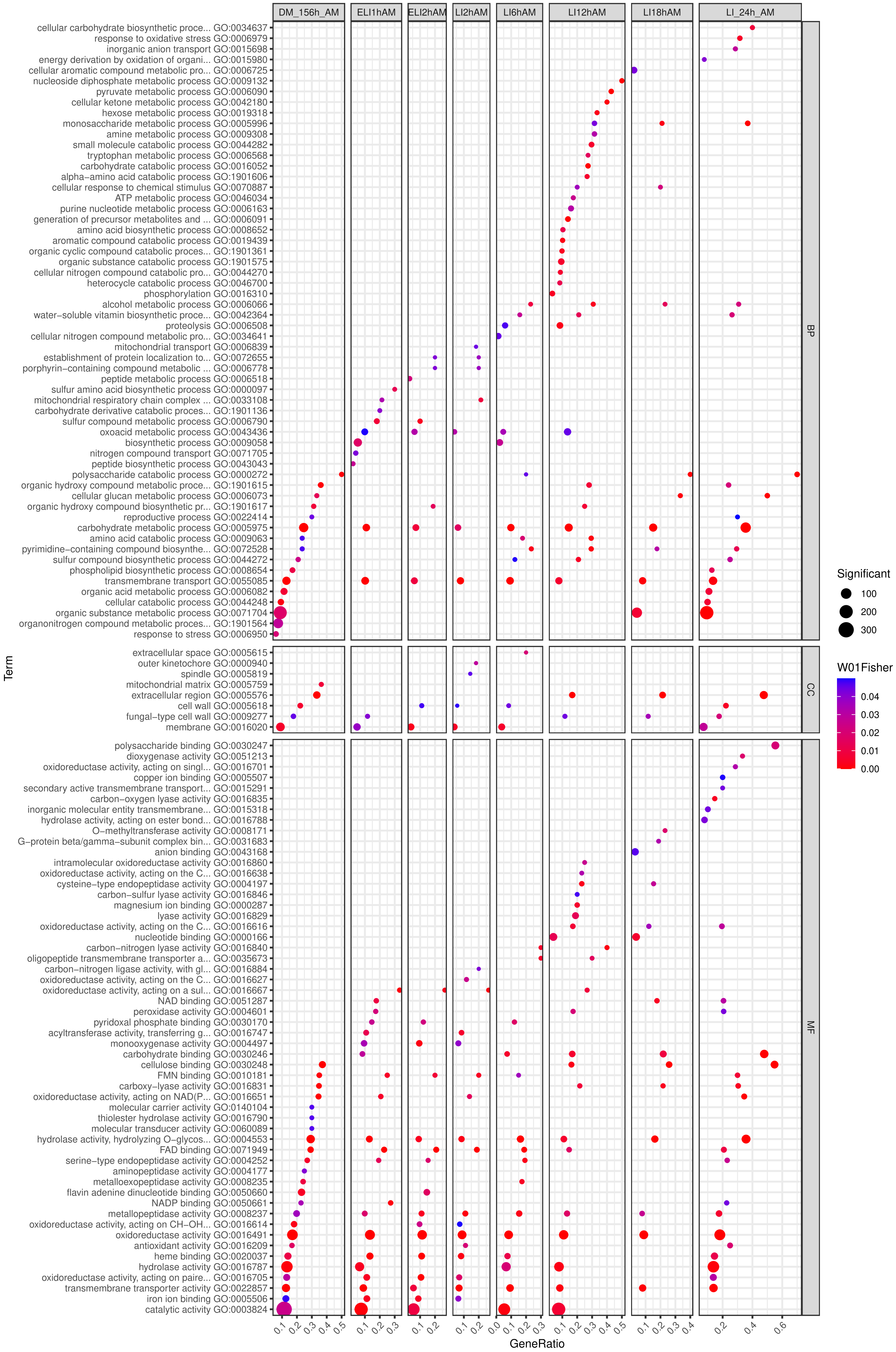


**Supplementary Figure 13.**
**Significantly enriched GO groups for light-induced attached mycelium (AM) samples.**

**Supplementary Figure 13.**

The y-axis represents the ratio of GO groups present in the gene sets. The size of the dots corresponds to the number of significantly expressed genes belonging to the GO group. The color of the dots indicates the statistical significance of the enrichment. The horizontal segmentation is based on the three main aspects of the Gene Ontology (GO) classification: Biological Process (BP), Cellular Component (CC), Molecular Function (MF).

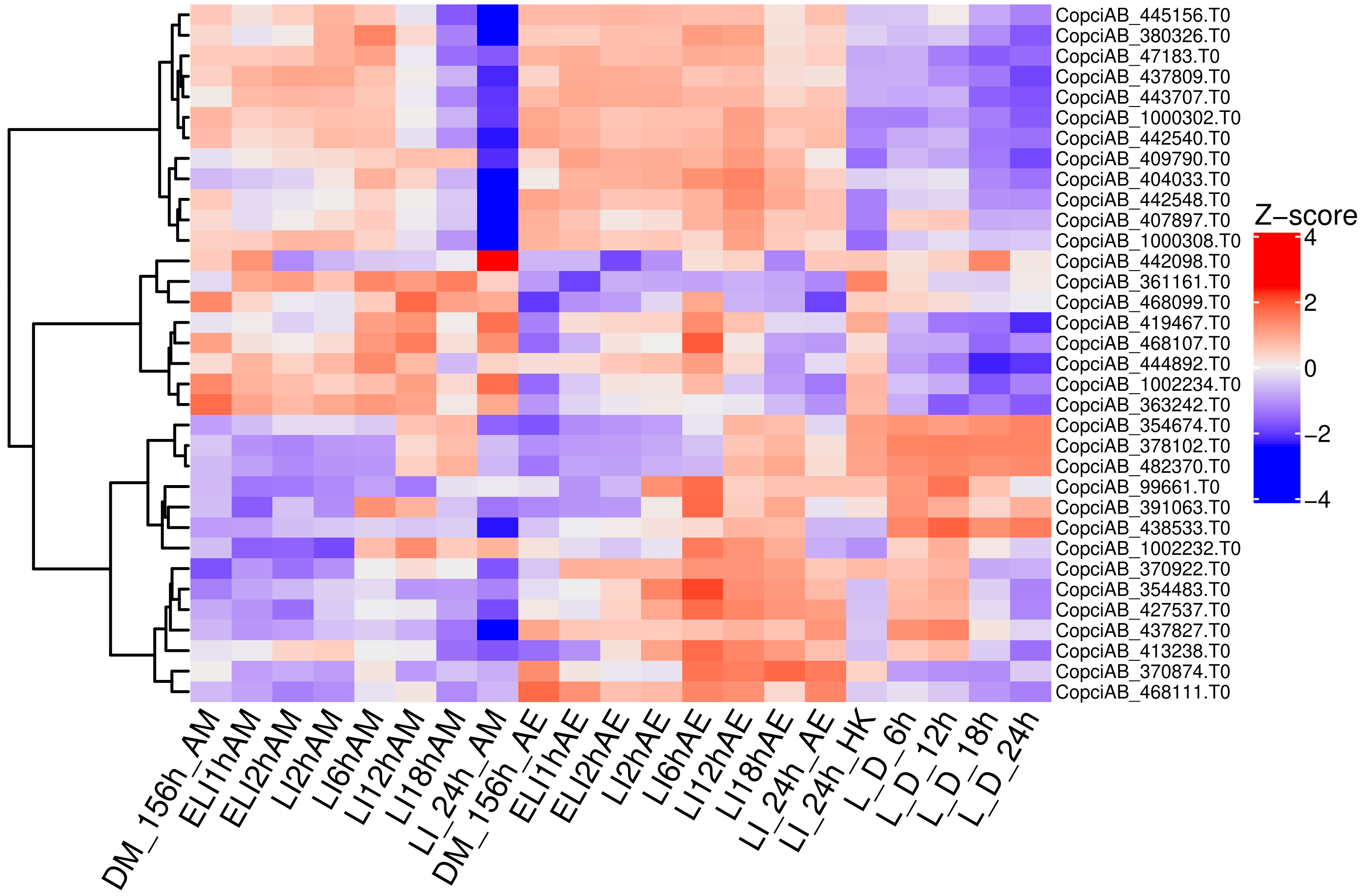


**Supplementary Figure 14.**

**Heatmap representation of the expression of the putative hydrophobin coding genes in the light induction related samples.** Gene expression values are represented by the mean $log_2$CPM of the biological replicants, which were scaled (Z-score) and hierarchically clustered by row (WardD method based on the Euclidean distance). In the case of sample names, 'DM', 'ELI', 'LI' and 'L_D' stands for dark grown mycelium, Early Light induction, Light induction and 12h Light/12h Dark cycle, respectively the length of incubation time in the given condition in hours. 'AE' and 'AM' stands for the aerial mycelium and attached mycelium.

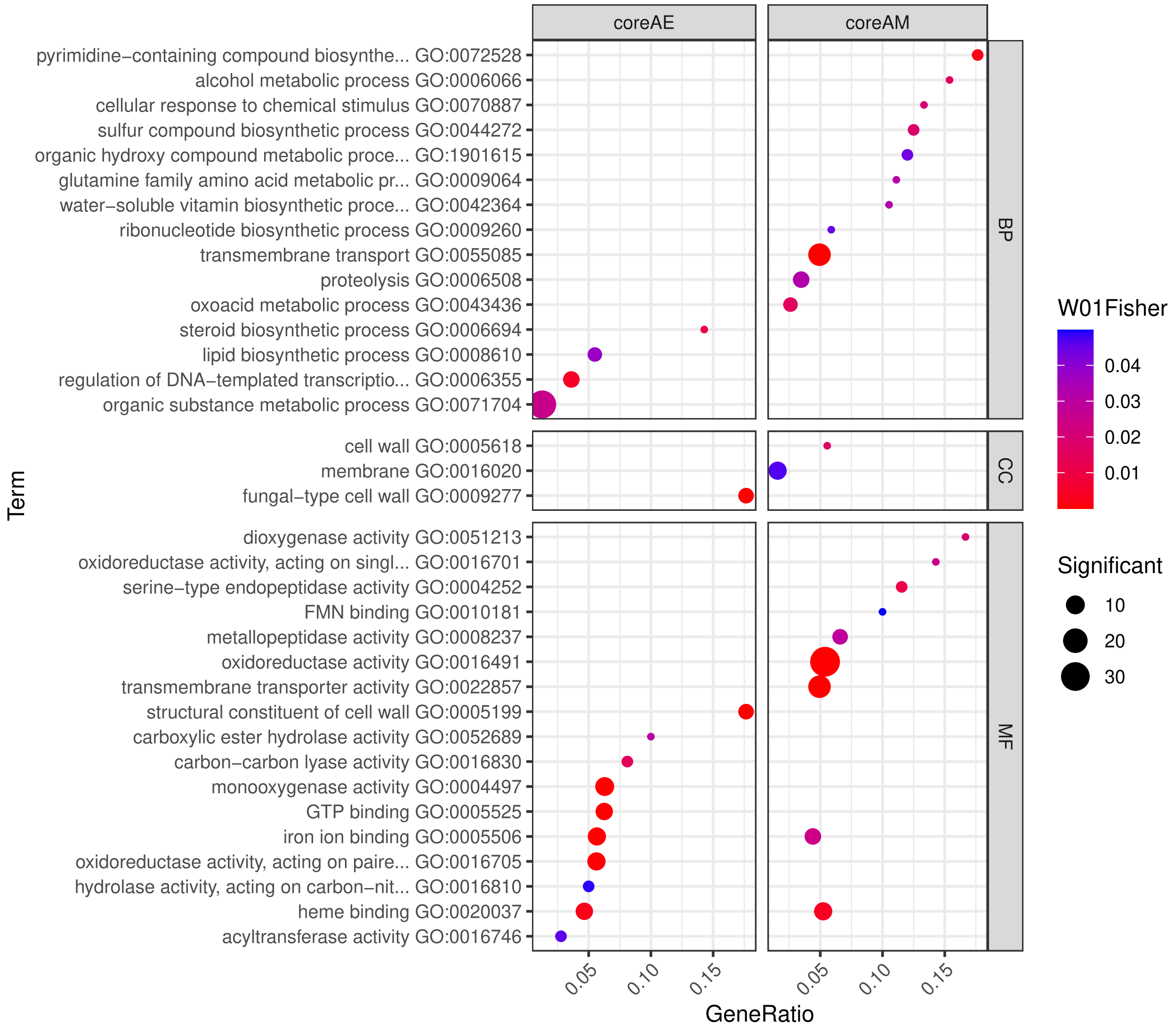


**Supplementary Figure 15.**

**Significantly enriched GO groups for core aerial and attached mycelium genes.** The y-axis represents the ratio of genes with a given GO present in the gene sets. The size of the dots corresponds to the number of significantly expressed genes belonging to the GO group. The color of the dots indicates the statistical significance of the enrichment. The horizontal segmentation is based on the three main aspects of the Gene Ontology (GO) classification: Biological Process (BP), Cellular Component (CC), Molecular Function (MF).
