## Supplementary Table 5 for "Unraveling Morphogenesis, Starvation, and Light Responses in a Mushroom-Forming Fungus, *Coprinopsis cinerea*, Using Long Read Sequencing and Extensive Expression Profiling"

**Development-related experiments:**

- **Basidiospore germination (BS) samples:** YMG/2 plates were inoculated with an agar cube and incubated in the dark for 5 days at 28°C. 2hrs of light treatment in plant chamber, then 24h in dark, then 12h/12h light/dark cycle. Spores collected after 5 or 6 days of light/dark cycle from fresh fruiting bodies in the morning. Spores that had fallen on the cellophane disk under the fruiting bodies have been collected. Cellophane disks were washed off in 0,1% Tween-20, filtered through a 40uM cell filter and centrifuged at 2700rpm for 10mins; pellet was washed with sterile DW once again with the same settings. Spores were diluted to $4x10^7$ and 250ul of this suspension was spread on YMG plates covered with cellophane disks and incubated at 37°C to induce spore germination. *Control sample in DEG analyses: BS_0h.*
- **Oidium germination samples**: For oidia production, YMG/2 agar plates were inoculated with a small agar cube and the colonies were grown in constant light at 37 °C. To harvest oidia, we poured 10 ml of distilled water on the surface of the colonies then gently scraped them with a blunt spatula. The suspension was then filtered through a 40 µm cell strainer (VWR). For oidia germination, 1 ml of oidia suspension ($10^8$/ml) was inoculated in 100 ml of liquid YMG in a 500 ml erlenmeyer flask and was incubated in the dark at 37 °C for 18 hours. *Control sample in DEG analyses: intact (non-germinating) oidia.*
- **Early light induction and light induction samples (ELI1h, ELI2h, LI2h, LI6h, LI12h, LI18h, LI24h):** We exposed YMG/2+cellophane 6.5 day old culture (grown in the dark at 28C) to white light for 2h. We sampled the edge of the colony only, with aerial and attached mycelium separately. Hyphal knots started to develop from 12h, these were impossible to separate from the mycelium samples, so they were sampled together. *Control sample in DEG analyses: DM_156h AM or AE.*
- **Light induction 24h (LI 24h) samples**: hyphal knots (just the mycelium with hyphal knots, as little amount of mycelium as possible), aerial mycelium without hyphal knots, and attached mycelium from between the middle of the plate and the hyphal knots.
- **Early fruiting in light/dark conditions (LD) samples**: Same as ELI samples, but following light induction, a 12h light/dark incubation at 28C was used. *Control sample in DEG analyses: LI_24h_HK*
- **Starvation response (SWAT samples):** place with cellophane, 2.5days old dark-incubated cultures from YMG/2 onto 1% water agar (in the dark) and sample all mycelium (areial + attached merged) 2, 4, 8, 16 and 24h. *Control sample in DEG analyses: DM_60h*
- **Hyphal knot formation in the dark (-30, -22,5, -15, -7,5, HK, HK + 7.5 samples):** 2,5 days old *C. cinerea* moved with cellophane from YMG/2 to 1% water agar in the dark incubated more at 28°C. 0,5 cm at the future HK part of the mycelium was sampled. *Control sample in DEG analyses: DM_60h.*
- **Sclerotium (Sclero samples):** YMG/2 plates grown at 28°C for 4 weeks in darkness. Sample the sclerotia scratched off as much as possible, with the least amount mycelia contamination possible. *Control sample in DEG analyses: DM_132h*

- **Dark Mycelium (DM) samples:** YMG/2 plates with cellophane disks were inoculated with an agar cube and incubated at 28°C in the dark for 12h, 36h, 60h, 84h, 108h, 132h or 156h. For all except the 156h sample the shole mycelium was sampled, for the 156h time point aerial and attached mycelia were sampled separately. *Control sample in DEG analyses: BS_12h.*
- **Cobalt chloride (Cob samples):** 2,5 days old *C. cinerea* moved in the dark from YMG/2 to 1% water agar containing 2mM cobalt-chloride. Sampled 30 hrs after WAT, -8 hours before HK should emerge. *Control sample in DEG analyses: CoK*

**Stress conditions, plated cultures:**

- **Frost (-20C):** YMG plates with cellophane grown at 28°C for 6 days in the dark, with the mycelium haven't touched the edge of the plate (Still growing). Plates covered in multiple layers of crumpled up paper placed for 30 minutes at -20°C. Whole mycelium was sampled. *Control sample in DEG analyses: DM_132h*
- **Heat shock (HS) samples:** YMG plates with cellophane grown at 28°C for 6 days in the dark, with the mycelium haven't touched the edge of the plate (Still growing) placed for 2h at 47°C. Whole mycelium collected in liquid N2. *Control sample in DEG analyses: DM_132h*
- **CO2 (CO2) samples:** YMG/2 plates with cellophane but plates sealed with electrical insulating tape airtight after inoculation grown at 28°C for 6 days in the dark, with the mycelium haven't touched the edge of the plate (Still growing). After the 6 days immediately after opening the plate the whole mycelium was collected in liquid N2. *Control sample in DEG analyses: DM_132h*
- **Drought (DR samples):** YMG/2 plates with cellophane grown at 28°C for 3 days in the dark (3 small cultures/ plate), with the cellophane transferred to YMG/2 containing 4% agar in the dark to avoid light induction. After incubation for 12h at 28°C we collected the whole mycelium. *Control sample in DEG analyses: DM_60h*
- **Trichoderma interaction (CTMP samples):** *C. cinerea* and *Trichoderma agressivum f. Agressivum (K9)* were inoculated at the same time on YMG/2+cellophane in the dark 28°C. At 2,5 days old we sampled only the mycelium at the edge of *C. cinerea,* where they met (not more than 6hours ago), sampled around 3 mm in only. *Control sample in DEG analyses: DM_60h*
- **Trichoderma interaction (CTGT samples):** *C. cinerea* and *Trichoderma* were inoculated at the same time on YMG/2+cellophane in the dark 28°C. They were grown together for 4,5 days, and the edge of *C. cinerea* mycelium (0,5-1cm) where they have been together the longest was sampled. *Control sample in DEG analyses: DM_108h*
- **Scratched mycelium (Scratchy samples):** YMG/2+cellophane 3.5 days old (still growing) *C. cinerea* culture. Cut the mycelium many times at the edge in the dark. Sample only the edge after 3 hours. *Control sample in DEG analyses: DM_80h*
- **Cold stimulation (Cold samples):** Shift 4 day old dark grown mycelium YMG/2 from 28C to 4C for 4 hours, then harvest. Whole mycelium was sampled. *Control sample in DEG analyses: DM_80h*
- **Hypoxia (Hyp samples)**: YMG/2 plates with cellophane 7 days old colony grown at 28°C. 4plates in BD Gaspack EZ hypoxia bag + 1 reagent pouch and 1 tissue paper

with 5ml of distilled water. Made small holes on the parafilm. Everything(transfer to gaspack) happened in the dark under red light. Covered them in aluminium foil and they were incubated for 24 hours at 28°C. Whole mycelium was sampled. *Control sample in DEG analyses: HypK*

- **Hypoxia control:** was treated the same but there were no reagent pouch and the bag was not sealed.

**Stress conditions, liquid cultures**: 250ml conical flask with 100ml of YMG broth grown at 28°C without shaking for 3 days. Only used the floating mycelia that had aerial and submerged parts as well. Cultures were stirred and disturbed but not teared due to the media modification. Control was always 'flask control'.

- **Flask control:** 3 days old mycelium in liquid YMG grown in the dark at 28°C - opened up, stirred - incubated for 12 more hours. Whole mycelium sampled. *Control sample in DEG analyses: LoKo*
- **Oxidative stress (H2O2 samples):** 1mM of $H_2O_2$ was added and incubated at 28°C for 3h. *Control sample in DEG analyses: LoKo*
- **Acidic culture (pH4 samples):** 3h incubation in pH set to 4 with HCl separately from the mycelium then poured back onto the mycelium. *Control sample in DEG analyses: LoKo*
- **Alkaline culture (pH9 samples):** 3h incubation in pH set to 9 with NaOH separately from the mycelium then poured back onto the mycelium. *Control sample in DEG analyses: LoKo*
- **Osmotic stress (Sor samples):** 10g Sorbitol/flask followed by 2h of incubation at 28C. *Control sample in DEG analyses: LoKo*
- **Cell wall stress (CR samples):** 1% Congo red added to the medium (Standard Fluka) and incubated for 3h. *Control sample in DEG analyses: LoKo*
- **Calcium ions (Calci samples):** Liquid minimal media without shaking growing in the dark. To 3 days old culture we added 200mM $Ca^{2+}$ for 24h, then the whole mycelium was sampled. *Control sample in DEG analyses: LoKo*
- **Copper ions (Cu samples):** Liquid minimal media without shaking growing in the dark. To 3 days old culture we added 2mM $CuCO_4$ for 6h, then the whole mycelium was sampled. *Control sample in DEG analyses: LoKo*
- **Voriconazole**: To 3 days old culture in YMG broth at 28°C we added 0.5ug/ml voriconazole (MIC 50 and 90) for 48 hours, then sampled the whole mycelium. *Control sample in DEG analyses: LoKo*
